# Pervasive within-host recombination and epistasis as major determinants of the molecular evolution of the Foot-and-Mouth Disease Virus capsid

**DOI:** 10.1101/271239

**Authors:** Luca Ferretti, Eva Pérez-Martín, Fuquan Zhang, François Maree, Lin-Mari de Klerk-Lorist, Louis van Schalkwykc, Nicholas D Juleff, Bryan Charleston, Paolo Ribeca

## Abstract

Although recombination is known to occur in FMDV, it is considered only a minor determinant of virus sequence diversity. This is because recombination appears to be highly suppressed at phylogenetic scales; inter-serotypic recombination events are rare; and in those a mosaic structure is present whereby recombination only occurs almost exclusively in non-structural proteins. Here we show that co-inoculation of closely related strains in buffaloes results over time in extensive within-host recombination in the genomic region coding for structural proteins. This enables us to directly estimate recombination rates for the first time. Quite surprisingly, the effective recombination rate in VP1 during the acute infection phase turns out to be about 0.1 per base per year, i.e. comparable to the mutation/substitution rate. Thanks to the features of our experimental setup, we are also able to build a high-resolution map of effective within-host recombination in the capsid-coding region. We find that the linkage disequilibrium pattern inside VP1 points to a mosaic structure with two main genetic blocks. Positive epistatic interactions between co-evolved variants appear to be present both within and between blocks. These interactions are due to intra-host selection both at the RNA and protein level. Overall our findings show that during FMDV co-infections by closely related strains, capsid-coding genes recombine within the host at a much higher rate than expected, despite the presence of strong constraints dictated by the capsid structure. Although those intra-host results are not immediately transportable to a phylogenetic setting, they force us to reconsider the relevance of recombination and epistasis, suggesting that they must play a major and so far underappreciated role in the molecular evolution of the virus at all time scales.

**Author summary:** Recombination in the capsid-coding region of the Foot-and-Mouth Disease virus genome is highly suppressed at phylogenetic scales. However, the role of recombination in the intra-host dynamics of the virus is not known. In our experiment, a co-infection of African buffaloes with closely related FMDV strains results in a population structure of the intra-host viral swarm, allowing us to detect recombination events. For structural protein-coding sequences, the swarm dynamics is driven by extensive within-host recombination. During the acute infection phase, we infer intra-host recombination rates of 0.1 per base per year, comparable to the typical mutation rate of the virus. The recombination map reveals two linkage blocks within the VP1 protein-coding sequence. Epistatic interactions between co-evolved mutations in VP1 are caused by intra-host selection at the RNA and protein level and are present both within and between blocks. Our findings support a major role for recombination and epistasis in the intra-host evolution of FMDV.

## Introduction

The foot-and-mouth disease virus (FMDV) is a picornavirus of the genus *Aphtovirus* that causes foot-and-mouth disease (FMD), an highly contagious vesicular disease. FMD is one of the most economically relevant diseases of livestock and cloven-hoofed animals [1] Domestic and wild artiodactyls usually develop viraemia a few days after exposure to FMDV, followed by an acute infection process that lasts about a week. In some cases such as African buffaloes, the infection progresses in a subclinical form and the virus can persist for years in carrier animals [2].

The FMDV genome is short (about 8000 nucleotides) and encodes a single ORF coding for a leader polypeptide (Lpro) that cleaves itself from the polyprotein, comprising four structural proteins (1A-1D or VP4, VP2, VP3, VP1) and nine non-structural proteins (2A-2C, 3A, 3B1-3B3, 3C, 3D) [3,4]. The determinants for infections and immunity are mostly found in VP1-VP4, which form the viral capsid.

Mutation rates in the FMDV genome are high, especially in the capsid-coding region. As it is often the case in RNA viruses [5], this is partly due to the lack of proof-reading capabilities of the polymerase. The high substitution rates contribute to the substantial genetic and antigenic variability of the virus. Seven different serotypes - A, O, C, Asia1 and Southern African Territories (SAT) 1/2/3 - are known, with a distribution spanning from south-eastern Asia to Africa and South America [6, 7]. SAT serotypes are endemic to Africa, where they circulate mostly among African buffaloes (*Syncerus caffer*).

Inside its animal hosts, the high mutation rates of FMDV may lead to the formation of a viral swarm, i.e. a cloud of similar genotypes differing only by a handful of mutations [8]. This is a typical pattern of intra-host genetic variability in organisms with high mutation rates, such as RNA viruses [5, 9] and is often correlated with a rich quasi-species dynamics [10].

Another important mechanism in the evolution of FMDV genomes is recombination [11–13]. Direct evidences of FMDV recombination date back 40 years ago [14,15]. Most recombination breakpoints are observed in non-structural proteins. Recombinant capsid-coding sequences have been described [16,17], but they appear to be much rarer than recombination events in non-structural proteins. More recently, systematic studies [18, 19] have found phylogenetic evidences of extensive recombination among non-structural proteins and only a small number of recombination events within capsid-coding sequences.

Recombination inferred from phylogenetic studies suffers from a strong detection bias. In fact, only events that occur between sufficiently divergent lineages can be detected, and only events that do not disrupt positive epistatic interactions among variants (i.e. events preserving correlated sets of genomic variants that taken together confer an evolutionary advantage to the virus) can generate viral sequences that are fit enough to be observed in samples [19, 20]. In addition, since the capsid proteins are the primary target of the immune response, cross-immunity of viruses with similar capsid-coding sequences could reduce co-infections and therefore recombination.

Within-host studies offer the opportunity to observe recombination in action without any of these biases [21,22]. Furthermore, intra-host recombination is an interesting subject in itself due to its role in the generation of genetic diversity within hosts [10,23,24]. In this respect, one of the best experimental systems for FMDV is arguably represented by infections in African buffalo, since animals of this species are FMDV carriers: after an initial acute phase of the infection, the virus can persist for years in some tissues, albeit at lower levels of replication [2]. In principle, this increases the chances to observe recombination events. The SAT serotypes of the virus are well-adapted to this host and there is evidence that buffaloes contribute to their dissemination [25].

In a recent experiment on African buffaloes infected by FMDV [2], viral sequences from different animals and tissues were generated with both Sanger and high-throughput sequencing technologies. An interesting feature of this experiment is the subsequent discovery of a strong genetic structure among the viral sequences. It turns out that both the inoculum and the animal samples contain two major viral swarms with moderate sequence divergence between them. Our results show that recombinants of these swarms were already present in the inoculum - probably due to previous recombination in buffalo or in culture - and the amount of recombination increased both after the acute phase of the infection and during the persistent phase. Thus this experimental system provides an excellent setup to infer the relative and absolute rates of within-host recombination.

In this paper, we present a detailed analysis of the genomic patterns of recombination in the capsid-coding region. First, we provide a brief explanation of the experimental setup and how it allows to detect within-host recombination. Then we present some estimates of the absolute recombination rates during the infection process - which turn out to be comparable to the mutation rates - and infer the recombination profile inside the capsid-coding region. Focusing on the VP1 coding region, we show how the linkage disequilibrium (LD) patterns among variants suggest a mosaic structure with two main blocks inside VP1. In addition, these patterns indicate the existence of intra-host epistasis between variants from different swarms, with epistatic interactions acting both within and between blocks. Finally, we discuss the evolutionary consequences of our findings both for the intra-host quasi-species dynamics and for the long-term evolution of FMDV, and explore the possible explanations behind the lack of recombination on phylogenetic scales.

## Results

### Co-infection and recombination of viral swarms

Details of the experiment are explained in [2]. In brief, a group of 16 buffaloes were co-infected with the three FMDV Southern African Territories serotypes (SAT1, SAT2, SAT3), and tissue samples were obtained at day 35 post-infection from two animals and day 400 from one animal. The inoculum contained equal amounts of SAT1, SAT2, and SAT3 virus, but only SAT1 was found in infected buffaloes one year after infection. The other serotypes play no known role in the dynamics of SAT1 diversity, hence in the rest of the paper we focus only on this serotype.

Interestingly, it turns out that the capsid-coding sequences of SAT1 viruses in the inoculum exhibit a strong multi-swarm structure, with most nucleotide sequences belonging to one of two major viral swarms (see Methods; according to the literature in the field previously mentioned, we use the term “viral swarm” to indicate a cloud of similar genotypes differing only by a few mutations). The VP1-coding sequences of the two swarms differ by about 3%, much larger than the genetic diversity within each swarm. Hence, the two swarms are clearly genetically distinct and separable. A similar multi-swarm structure is found in viral VP1-coding sequences from micro-dissections of several tissues from three infected buffaloes, proving that co-infection occurred in this experiment. The genetic variability in these sequences is illustrated in Figure 1. A detailed analysis of these swarms, their evolution post inoculation and the immune response of the animals will be presented elsewhere (Cortey et al, companion paper).

**Fig 1.**
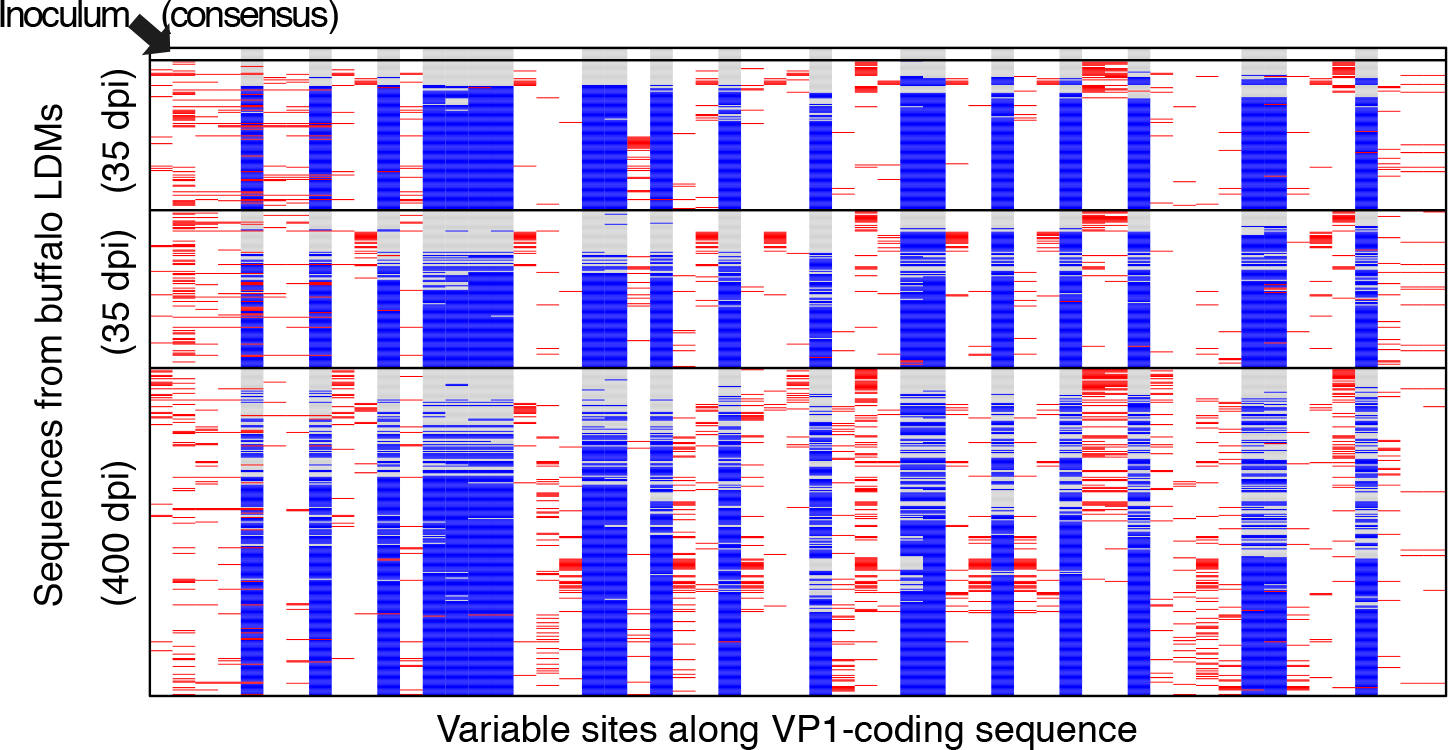
Illustration of the content of the VP1-coding sequences from buffalo tissues. Sequences are sorted by animal, then by divergence from the consensus sequence of the inoculum. New mutations with respect to the consensus sequences of the two swarms are in red, while inter-swarm mutations characterising the two initial swarms are in grey and blue, respectively.

Co-infection of these buffalo hosts by different swarms offers an opportunity to observe within-host recombination. In fact, recombination is assumed to occur whenever two viruses co-infect the same cell [20], but it can only be detected when their sequences are different enough to be clearly separated. This is the case in our experiment. Indeed we observe a large number of recombinants among the Sanger sequences of clones derived from the buffalo tissue micro-dissections (Figure 1). Extensive recombination between sequences belonging to the two initial swarms is also detected in the short reads from the inoculum. These observations cannot be explained by sequencing errors, chimeric artefacts or repeated mutations (see Methods).

The amount of recombination in these sequences is surprisingly large: there is at least one recombination event between almost all pairs of Single Nucleotide Variants (SNVs) characterising the two swarms. Furthermore, the fraction of recombinants seems to increase in time post inoculation (Figure 2), suggesting that the mixture of co-infecting swarms has a recent origin and has not reached a stationary equilibrium. These features allow us to apply classical population genetics approaches to this system.

**Fig 2.**
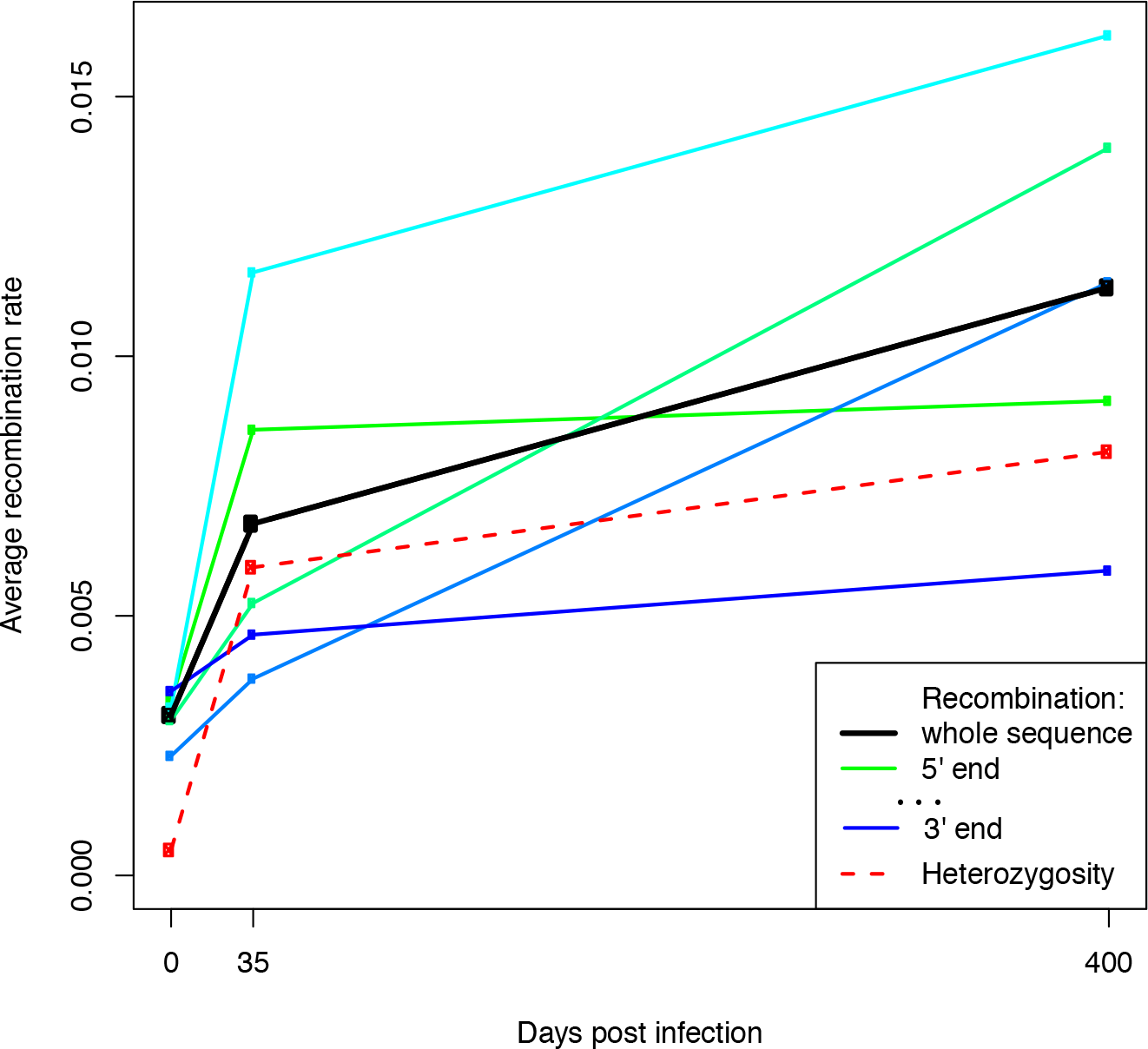
Cumulative recombination rates in VP1-coding sequences from the inoculum and from animals sampled at 35 and 400 dpi. All the rates are defined from the beginning of the experiment to the sampling time of the sequences and are computed using the “local” approach. Different colours illustrate the behaviour of different non-overlapping portions of the VP1-coding sequence, with a green-blue gradient from the 5’ to the 3’ end. The dashed red line shows the heterozygosity per base, computed on the intra-swarm SNVs (i.e. variants unrelated to the main swarm structure).

In classical population genetics, recombination can be inferred from *linkage disequilibrium* (LD), a measure of the correspondence between the genotypes of two closely occurring SNVs [26]. In the absence of recombination (and of recurrent mutations), the physical linkage between alleles along the sequence constrains the possible allelic combinations. As an example, for two SNVs originating from a mutation ...A...G...→...T...G... in the first site followed by a ...A...G...→...A...C... mutation in the second site, the only possible allelic combinations without recombination are {A,G}, {T,G} and {A,C}, i.e. C in the second site would always be found with A in the first, and T in the first site would always be found with G in the second; in addition, {A,C} would tend to appear at lower frequencies. The effect of recombination events between the two SNVs is to reshuffle these allelic combinations; in our example, recombination occurring between the two SNVs would generate sequences with a {T,C} genotype and increase the frequency of the {A,C} genotype. Linkage disequilibrium quantifies the observed extent of reshuffling between genotypes.

There is a direct relation between the strength of LD and the recombination rates. If the sequence between two SNVs recombines at a rate *r* for a time *t* since the formation of the mixture, LD decays over time as an exponential LD ~ *e*^−*R*^ of the cumulative recombination rate *R* = *r* · *t* [26]. However, linkage disequilibrium could also be affected by *epistasis*, i.e. fitness-related interactions between genetic variants. In fact, when recombination disrupts favourable combinations of co-evolved variants, recombinants have lower fitness and their number is suppressed by selection. More generally, if different combinations of alleles at multiple loci have different fitness, the frequency of favoured combinations of alleles increases and the synergy between alleles corresponding to these combinations is reinforced. Hence these epistatic interactions often act in opposition to recombination and cause an effective increase in LD [27].

### Absolute recombination rates

Linkage disequilibrium and recombination rates were inferred for the whole capsid-coding region of the inoculum and for the VP1-coding sequence of the virus from three buffaloes, two sampled at 35 days post infection (dpi) and one sampled at 400 dpi. All cumulative recombination rates inferred from our sequence data are relative to the time of origin of the mixture of swarms, which is not known, hence their absolute values do not have any easy interpretation. However, their differences provide absolute recombination rates per unit time across the acute and persistent phases of the infection. We can estimate these rates only for VP1, since it is the only genomic region for which multiple time-points are available.

Recombination rates were estimated using LD between pairs of inter-swarm SNVs, i.e. variants consistent with the two main swarms of the inoculum. Two approaches were used for inference of recombination rates: the “local” approach uses only information from consecutive variants, while the “global” approach uses information from all variants. The “local” approach is therefore more noisy, while the “global” one is more precise but could be more sensitive to biases. The two methods are also affected by epistatic interactions, but at different scales.

The average cumulative recombination rates per base, estimated using the “global” and “local” approaches, are *R*_0_ ≈ 2.6 · 10^−3^ − 3.0 · 10^−3^, *R*_35_ ≈ 4 · 10^−3^ − 7 · 10^−3^ and R400 ~ 8.0 · 10^−3^ − 11.7 · 10^−3^ respectively, accumulating in time as illustrated in Figure 2. Hence, the rate per year during the first 35 days post inoculation is *r*_0−35_ ≈ 0.015 − 0.040/site/y, while for later times the rate is *r*_35-400_ ≈ 0.004 − 0.005/site/y. Hence, during the first month post-infection, the average recombination rate is higher by a factor *r*_0−35_/*r*_35−400_ ≈ 3.8 − 8.7. Since the acute phase of the infection lasts about a week [2], the actual rates from the “global” and “local” approach can be estimated as

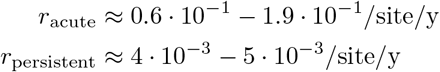

i.e. recombination during the acute phase is 15 - 40 times faster than during the persistent phase.

Note that these absolute recombination rates are comparable or even higher than the typical mutation rates for FMDV, which are as high as 10^−2^ mutations per base per year [28,29] due to the error-prone nature of the RNA polymerase. The rates per site per generation are also of the same order of magnitude as the ones inferred for *in vivo* HIV infections [23].

### Recombination profile in the capsid-coding region

We now look at the fine-scale structure of recombination rates. The basis for the inference of recombination is the normalised linkage disequilibrium *D*′ between pairs of derived swarm-specific variants. The measure *D*′ is defined in the Methods and it takes values +1 or −1 in the absence of recombination, while it is close to 0 for strong recombination. The *D*′ values for pairs of variants in the capsid-coding region are shown in Figure S2 as estimated from high-throughput reads from the inoculum.

A detailed recombination profile can then be built from *D*′ using the “global” and “local” approaches discussed in the previous section. This recombination map extends almost to the whole sequenced region, i.e. capsid-coding plus flanking regions, and the distance between swarm-specific variants (~ 30 bases on average) determines the resolution of the profile. The final profile is shown in Figure 3.

We observe that recombination rates inferred by the “local” approach for the capsid-coding region peak strongly around the 3’ end of Lpro/5’ end of VP4. They also show a moderate heterogeneity both between and within protein-coding regions, with peaks around the middle of the VP4- and VP3-coding sequence and in the region of 2A-B.

**Fig 3.**
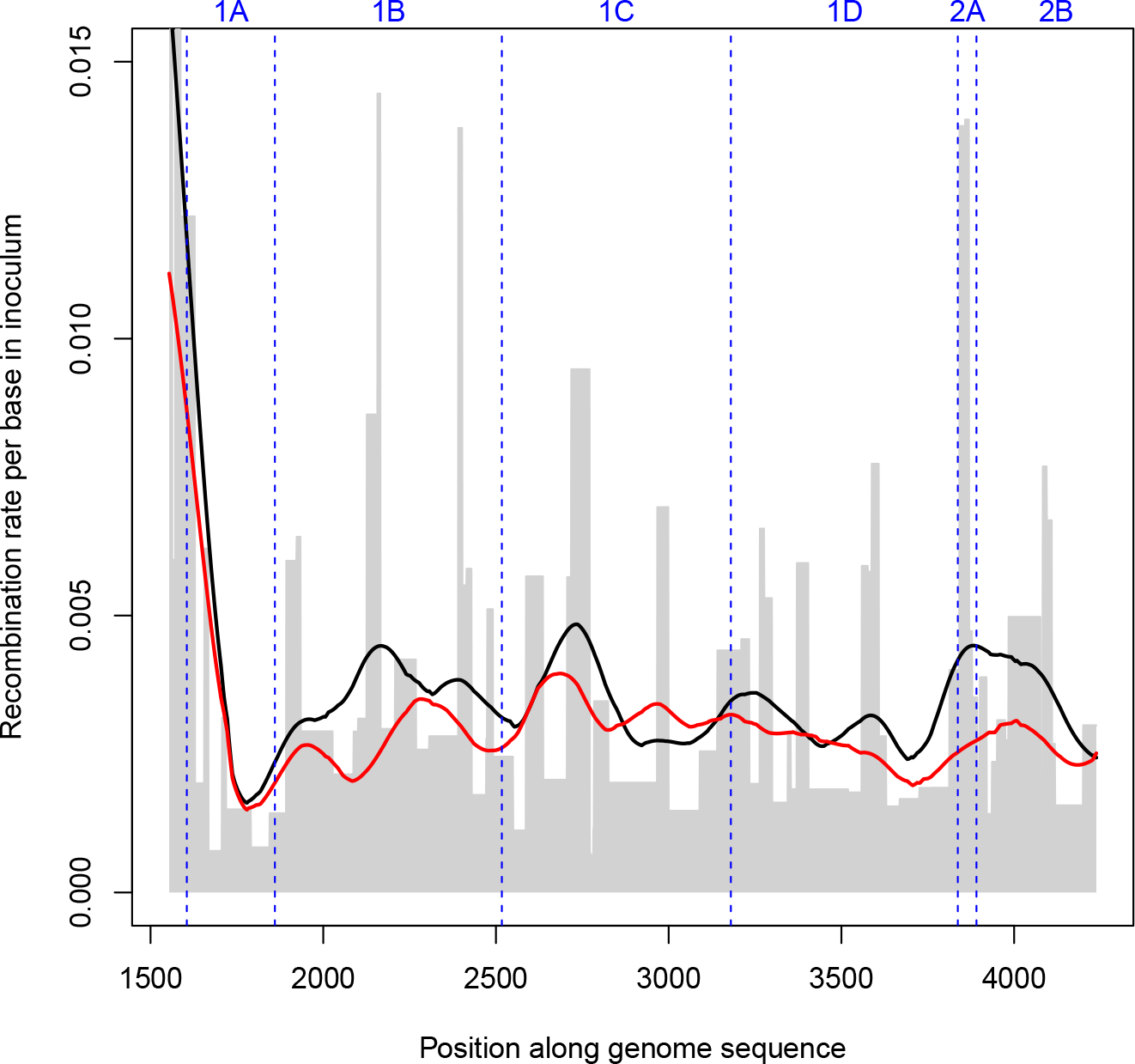
Effective cumulative recombination rate per base for the inoculum (in grey), inferred by the “local” approach from pairs of SNPs covered by at least 10^3^ reads. The lines indicate the average local rates (black) and global rates (red) Gaussian-smoothed over a 150 bases window.

From this recombination map, it is also possible to obtain an estimate of the relative recombination rates with respect to the VP1-coding sequence of the other capsid protein-coding sequences (VP4, VP2, VP3 or 1A-1C) and some non-structural protein-coding ones (2A-2B and small regions at the 5’ end of 2C and at the 3’-end of Lpro). These relative rates are summarised in the following table:

| Region | $R/R_{VP1}$, global | $R/R_{VP1}$, local |
| --- | --- | --- |
| Lpro | 6.2 | 8.7 |
| 2A | 1.12 | 2.8 |
| 2B-2C | 1.04 | 1.17 |
| 2A-2C | 1.05 | 1.4 |
| VP4 (1A) | 0.93 | 0.87 |
| VP2 (1B) | 1.07 | 1.27 |
| VP3 (1C) | 1.26 | 1.15 |
| 1A-1C | 1.12 | 1.16 |

Recombination rates in flanking regions of the capsid-coding sequence (Lpro and 2A) are higher than in the capsid-coding sequence itself. Hotspots of recombination in the flanking regions of the capsid-coding sequence have been previously detected in studies based on phylogenetic evidence [18,19]. These previous analyses inferred levels of phylogenetic recombination that were extremely low for VP1- and capsid-coding sequences and much higher for non-structural protein-coding ones. In partial contrast with these studies, we observe high intra-host recombination rates in the capsid-coding region, while the recombination rate in 2A is larger but still of the same order of magnitude as the capsid rate, and the rates in 2B-2C and in the capsid-coding region are actually similar.

### Mosaic structure and epistasis in the VP1-coding region

Thanks to our experimental design, recombination profiles for the sequence coding for VP1 (1D) can be reconstructed from different individuals and timepoints: the inoculum, two animals sampled at 35 dpi and an animal sampled at 400 dpi. It is therefore interesting to compare the different profiles. The absolute recombination rates inferred from the “local” approach are shown in Figure 4. On the top of a trend of increasing recombination rates with time (already clear in Figure 2), we observe some heterogeneity in the recombination rates along the protein-coding sequence, with several peaks found in similar locations across different individuals.

**Fig 4.**
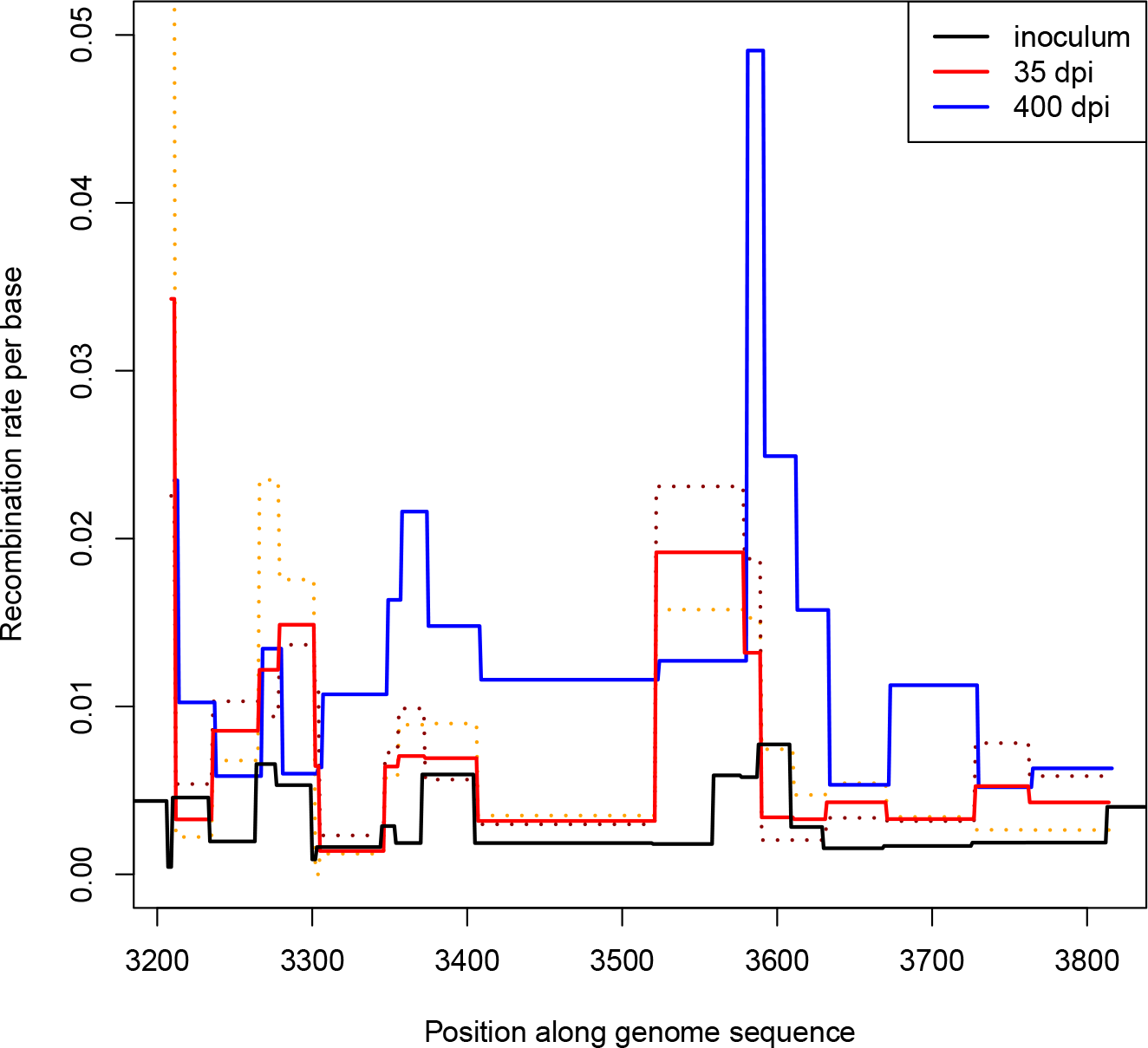
Effective recombination rate per base along the VP1-coding sequence. The cumulative rates are measured from the beginning of the experiment to sampling times: 0 days post inoculation (dpi) i.e. inoculum; 35 dpi; and 400 dpi. To illustrate the heterogeneity in inferred recombination rates between individuals, two separate dashed lines are shown for the two individuals sampled at 35 dpi.

The complete VP1-coding sequences from micro-dissections of buffalo tissues reveal a richer structure created by the interplay of recombination and epistasis. In fact, these sequences contain information about the linkage disequilibrium of most pairs of variants within VP1, as they have been obtained by Sanger sequencing. The corresponding LD maps for inter-swarm SNVs are shown in Figures 5A-D (lower triangles). The LD patterns show two strongly linked blocks with |*D*′| ≳ 0.5, roughly corresponding to the first 200 and last 250 bases of the VP1-coding sequence. These blocks correspond to the red-orange triangles in the lower half of Figure 5A, and their pattern is broadly consistent across different individuals and times (Figures 5B-D). This suggests that the mosaic structure observed in the non-structural part of FMDV genomes [19] is not restricted to non-structural proteins, but actually extends to the capsid-coding region. Recombination interacts with other forces within the host to maintain a modular structure with at least two different linked genetic blocks inside the VP1 protein-coding sequence.

**Fig 5.**
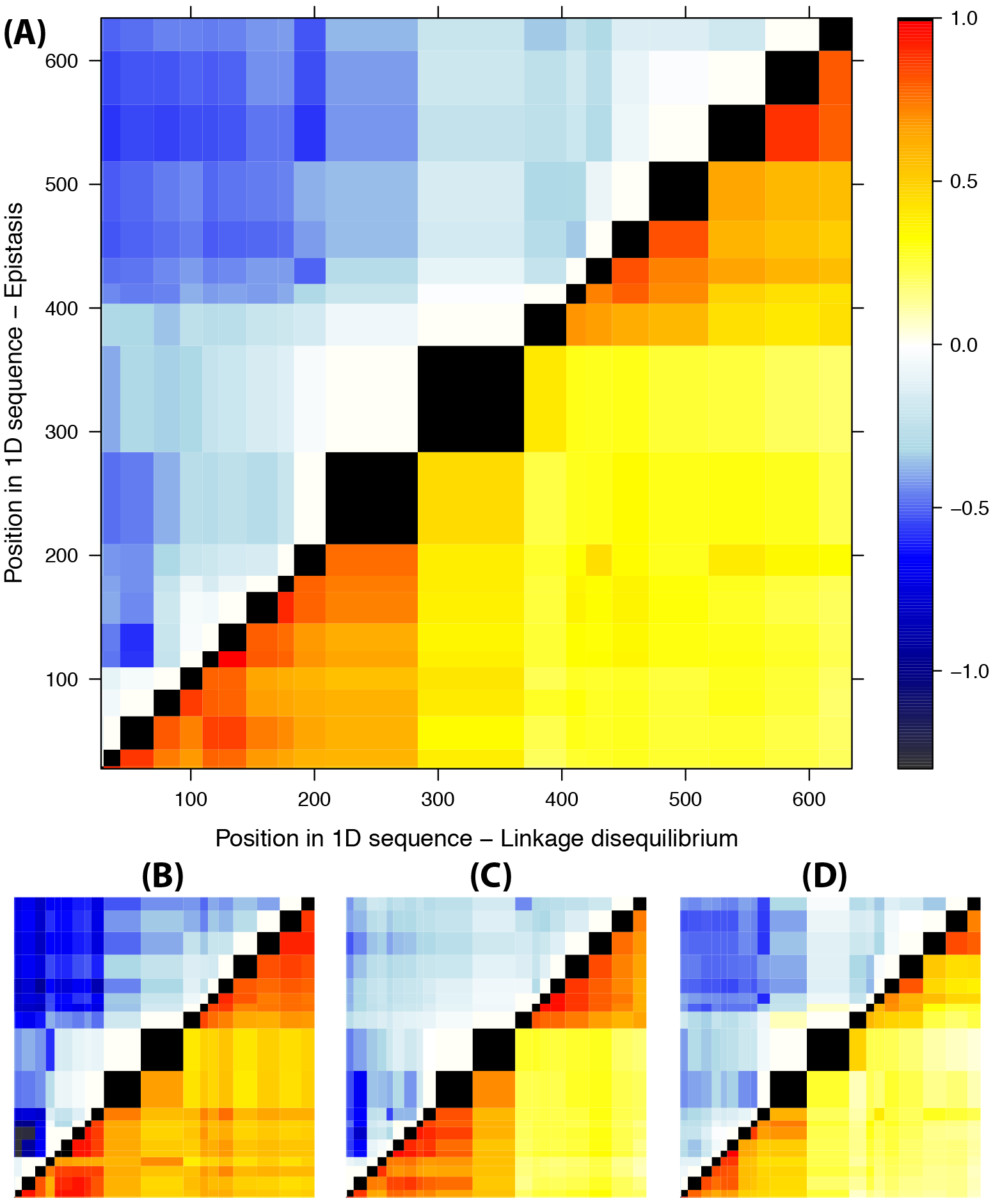
LD and epistasis across sequences from all animals (A), the two individuals sampled at 35 dpi (B,C) and the individual sampled at 400 dpi (D). Lower triangles: map of pairwise LD between variants in VP1, estimated as |*D*′| (stronger LD shown in red). Upper triangles: signatures of epistasis in terms of effective suppression of recombination log_10_(*R*/*R*_predicted_) (stronger epistasis shown in blue).

Epistasis is another major force influencing and possibly driving the intra-host dynamics. The LD between variants in the same protein-coding region can be affected by epistatic interactions due to functional constraints, stability or immune pressure. If the original combinations of variants in the swarms are fitter than the recombinants, the recombination rate is effectively reduced by selection [26,27]. The effects of recombination rate and epistasis cannot be separated for pairs of consecutive variants, since it corresponds to the scale of the finest resolution of LD, and the only information available at this scale is a single measure of *D*′. However, for distant pairs of variants, it is possible to detect footprints of epistatic interactions from excess of LD with respect to the naive value 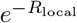 estimated from the “local” approach to recombination rates. We can use the effective suppression of recombination *R*/*R*_local_ as a signature of epistatic effects. We also infer the strength of selection against recombinants based on an explicit model of evolution with pairwise epistatic interactions (see Supplementary Information). The strength of the epistatic selection coefficients leads to qualitatively similar results as the above measure of suppression of recombination.

As expected, intra-protein epistatic interactions shape the LD structure of the VP1-coding region. In fact, the suppression of recombination in Figures 5A-D (upper triangles) hints at the presence of epistatic interactions inside both genomic blocks in the VP1-coding region. These interactions could contribute to its modular structure [30]. Interestingly, strong signatures of epistasis are found between the two blocks as well. This indicates that even if recombination tends to decouple the two blocks, linkage equilibrium is prevented by epistatic interactions between the blocks, which suppress replication and infectivity of recombinant sequences.

Given the large number of interacting variants, it is difficult to disentangle the strength of each pairwise interaction from the cooperative epistatic effects of all other linked variants; their cumulative effect could lead to “genotype selection”, i.e. locking variants into haplotypes containing only the most favourable combinations [30]. We modify the local prediction of recombination rates to a nonlocal one *R*_nonlocal_ that accounts for the suppression due to the most strongly linked variant between each pair.

With this finer approach, the observed suppression of recombination *R*/*R*_nonlocal_ confirms the presence of stronger interactions within the two blocks and weaker ones between blocks. We also observe that the recombination tends to be more suppressed between non-synonymous variants (Figure S4). In fact, the strength of epistatic interactions between pairs of non-synonymous variants is significantly higher than between other pairs at similar distances (*p* = 0.04, Figure S5). This proves that the selection pressure acts at the protein level as well. Selection on the aminoacid sequence is likely to be related to the stability of the VP1 protein structure, as apparent from the exposure of interacting aminoacids (3 out of 5 are buried), the lack of aminoacids in known epitopes (only 1 out of 5) and the spatial proximity of the interacting aminoacid pairs in the capsid structure (Figure 6).

**FIG 6.**
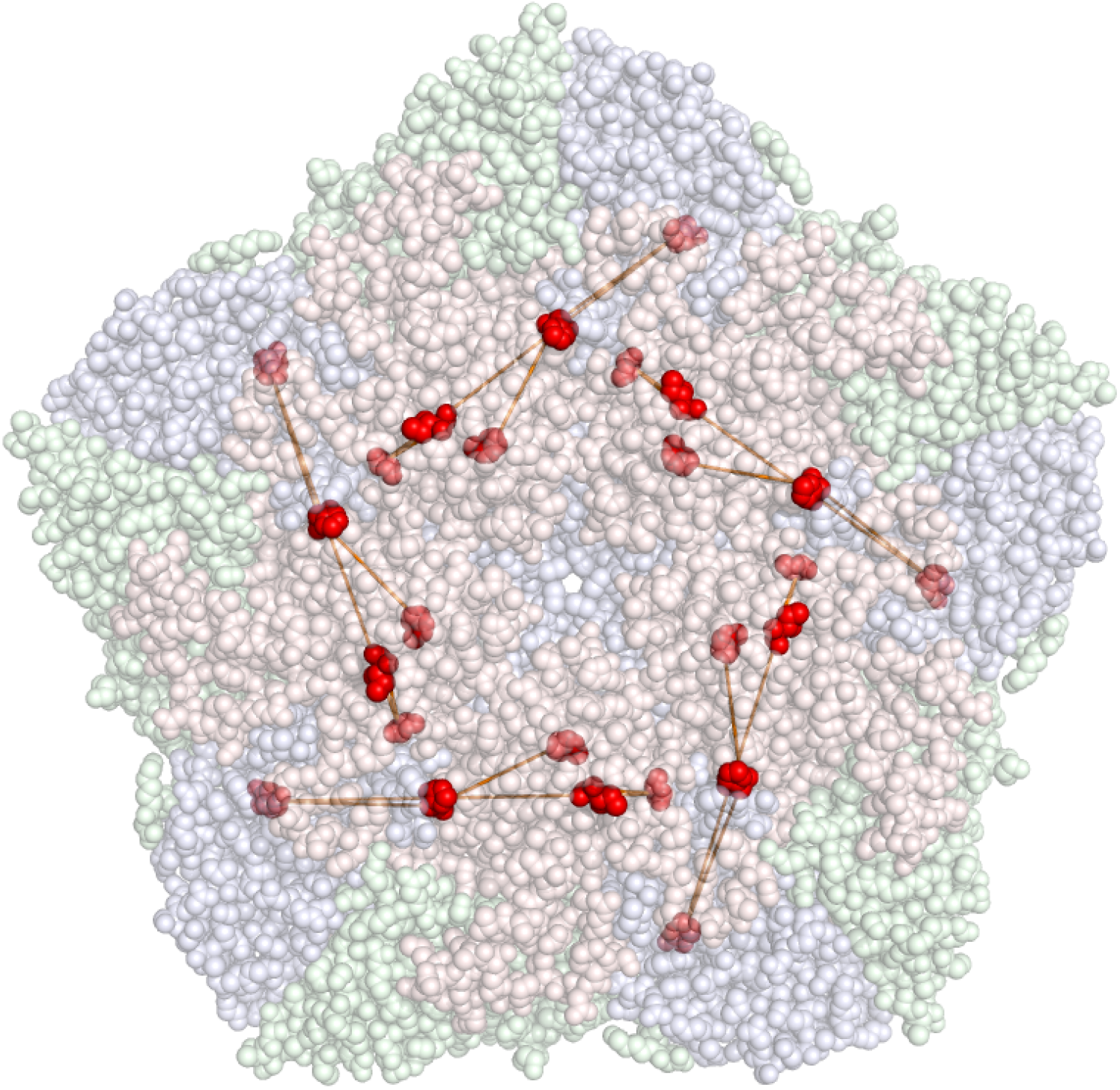
Localisation of the aminoacids corresponding to the five inter-swarm non-synonymous variants involved in epistatic interactions, projected on the capsid structure of O/BFS-1860. The red, green and blue components of the capsid correspond to VP1, VP2 and VP3 respectively. Variants corresponding to exposed aminoacids are shown in darker colors.

Overall, these results imply that epistatic interactions are widespread inside the VP1-coding region. Intra-host selection acts both at the RNA and protein level and epistatic interactions exist at both levels. In our experiment, these interactions reduce the rate of recombination between blocks by up to an order of magnitude, corresponding to selection coefficients of order 10^−2^ between pairs of variants.

## Discussion

### Intra-host recombination map for the capsid sequence

The recombination maps both for the whole capsid-coding region and for VP1 have been inferred for infections of a single serotype (SAT1) in a single host (buffalo). The details of the recombination map might differ between serotypes. The absolute rates and the amount of epistasis are expected to depend on the within-host infection dynamics and the immune response of the host, hence they could vary as well when considering infections in cattle or other species. However, we do not expect the qualitative picture for other host species and serotypes to be essentially different.

The inferred recombination rates are related to the amount of co-infections of the same cells in the host, hence they depend intrinsically on within-host dynamics [20]. This is unavoidable in all studies of viral recombination in vivo. In fact, it is better to consider our results as “effective recombination rates” which already take into account the effect of within-host evolution and infection dynamics. It might be possible to study recombination in a more controlled setup: in vitro and in vivo co-infection experiments guided by our findings represent a promising avenue to a more systematic inference of recombination rates.

#### Contribution of epistasis

Recombination rates depend on epistasis as well. This is unavoidable even when considering high-resolution maps such as ours, since we have no way to account for the effect of local epistatic interactions between close mutations. However, we were able to detect intra-host epistasis on the scale of the VP1 sequence and discovered that epistatic interactions are widespread. This also means that “local” estimates, which are affected only by local epistasis, should be more reliable than “global” estimates, which are more likely to systematically underestimate the real rates due to the additional effect of longer-range epistasis and the cumulative effect of cooperative interactions among multiple linked variants. In fact, such underestimation appears clearly in Figure 3. This is also the reason why we presented only “local” estimates for most of our results.

### Consequences for the evolution of the FMDV capsid

The high recombination rates in structural proteins between genetically close lineages represent an important finding of this work. In fact, it is natural to assume that recombination between genetically closer sequences will be even higher. This has potentially relevant implications for the genetic diversity in quasi-species. In fact, mutation and recombination play different roles in generating genetic diversity, and their balance can affect the fate of the quasi-species, as recently suggested in [31]. Mutations have a direct effect on the diversity of the swarm by generating new nucleotide variants, but a high mutation rate also adds a significant load to the population, as most of these polymorphisms are deleterious and tend to reduce the overall fitness of the quasi-species [9]. On the other hand, recombination has an indirect role, by generating different combinations of existing nucleotide variants. This increases the diversity of the swarm while unlinking the fate of potentially advantageous and deleterious mutations increasing the chances of compensatory combinations of mutations, and reducing the probability of fixation of deleterious mutations [31,32]. All these effects alleviate the mutational load. Hence, although the actual role of recombination in RNA viruses is still unclear [33], high intra-host recombination rate could be highly beneficial for FMDV quasi-species and play an important role in their pathogenicity [20].

Recombination events could even generate new genetic diversity at a phylogenetic level in sequences coding for capsid proteins, provided that they are not suppressed by lineage competition or epistatic selection against recombinants. Further studies are needed to understand which phenomena suppress capsid recombination on broad epidemiological scales and which viruses with a recombinant capsid-coding sequence could represent an epidemiological risk.

We will discuss the most interesting evolutionary consequences of high recombination rates and epistasis in the capsid sequence in more details in the next sections.

#### Recombination as a leading force in generating within-host diversity

One of the features of RNA viruses such as FMDV is the formation of viral swarms and quasi-species. This is a consequence of the high mutation rate and the dependence of pathogenicity on the genetic diversity of the quasi-species [10]. The actual amount of genetic diversity within the swarm depends on the length of the infection, on the transmission bottlenecks and on the equilibrium between mutation, recombination and the selective pressures on the quasi-species. Genetic diversity in quasi-species has functional roles that are not fully understood: among others, it increases pathogenicity and adaptation to specific tissues [9,10].

The rates of intra-host recombination observed in this experiment suggest that recombination could be a leading force in generating genetic diversity in the swarms. In fact, a simple calculation using the substitution rates known from the literature shows that in an infected buffalo only about half of the FMDV sequences at the end of the acute infection phase would differ from the inoculum, most of them by a single nucleotide mutation; on the other hand, according to the rates observed here, most sequences would be recombinants of 4-5 viruses in the swarm. Most of these recombination events would be between almost identical viruses and do not lead to new haplotypes, but a sizeable fraction of events would create new haplotype combinations separated from the inoculum by multiple mutations, therefore enhancing the breadth of the swarm in genotype space while alleviating the mutational load by disentangling the fate of beneficial and deleterious mutations [31]. This effect is illustrated in Fig 7A.

**Fig 7.**
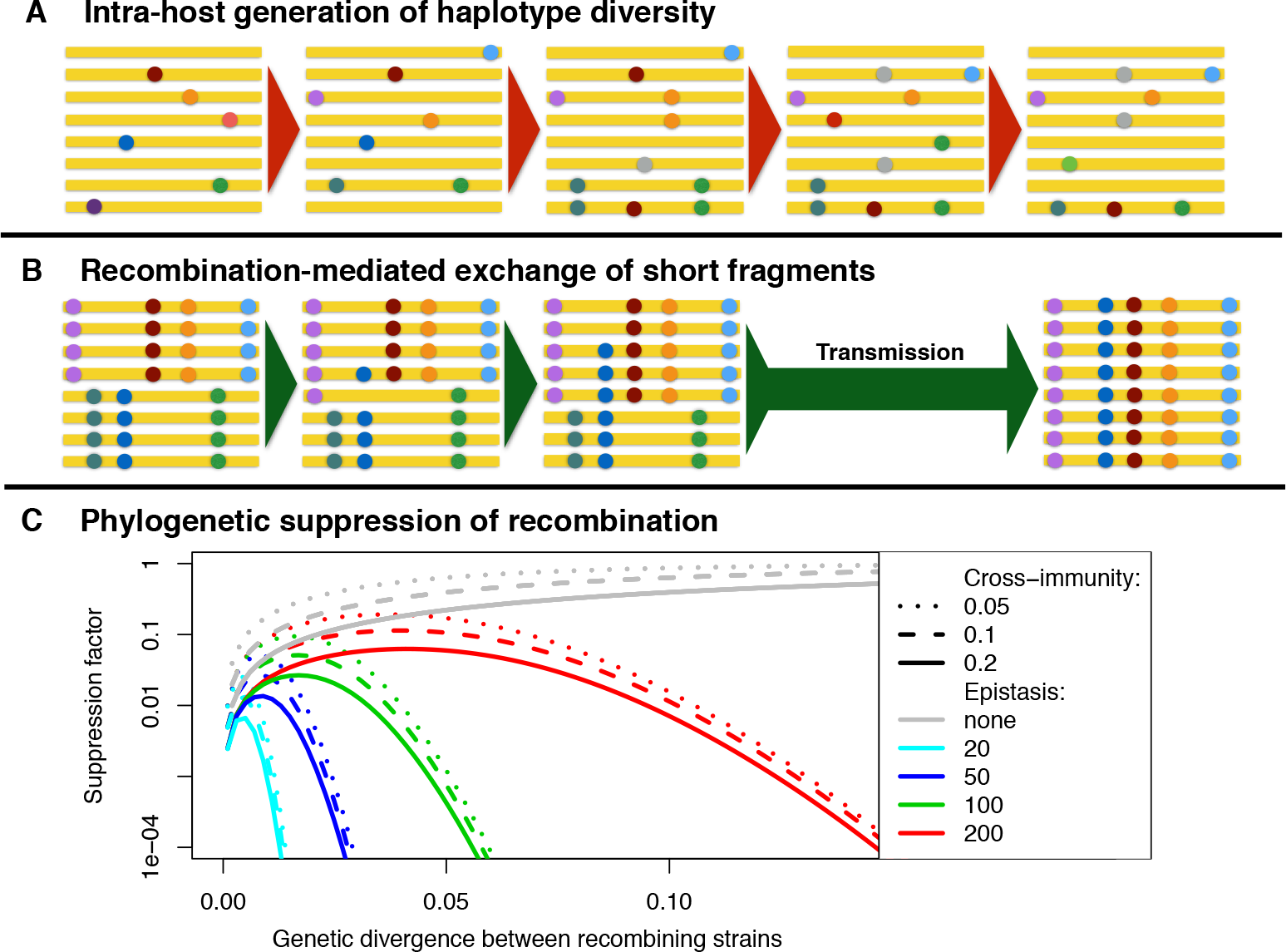
Illustration of the evolutionary consequences of recombination. A:Evolution of a swarm viral sequences (in yellow) and variants therein (coloured dots) under mutation and recombination. Intra-host recombination cannot generate new mutations in the swarm, but it generates new haplotypes from existing mutations and it disentangles the evolutionary fate of different mutations, reducing the effect of deleterious mutations and the mutational load. B: During co-infections by divergent strains, recombination may cause the exchange of short sequence fragments between strains. If the resulting viral strains are similar to the original ones, they may be transmitted despite the selective pressures against recombinants, as illustrated. C: Plot of the predicted reduction in combination in structural proteins at phylogenetic scales, due to cross-immunity and epistatic interactions, shown as a function of the genetic divergence (i.e. Hamming distance per base) between recombining sequences. Cross-immunity is modelled as an exponentially decreasing function of the divergence *d* between strains, following an approach used for influenza [47]. Assuming that cross-immunity acts between lineages with divergence less than *d*_*ci*_, the reduction in recombination corresponds to the reduction in co-infection 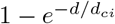. Epistasis is modelled after [43] as a decrease in viral growth rate proportional to the number of pairwise interactions disrupted by recombination, leading to a recombination suppression factor 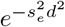 where *d* is the genetic divergence and the epistatic coefficient *s*_*e*_ is related to the strength and number of epistatic interactions. Assuming that the strength of epistasis among different capsid protein-coding sequences is is comparable to the one observed within host between the two blocks of VP1, an approximate estimate for this coefficient could be around *s*_*e*_ ~ 50.

The multi-swarm structure of our experiment offers some indirect evidence to back these observations. We perform a further analysis on the viral sequences from buffalo tissues, ignoring the intermediate-frequency SNVs that distinguish the two swarms, and focus on the low-frequency SNVs within each swarm. The corresponding genetic diversity is a good representation of the potential intra-host diversity of a single quasi-species. For each buffalo, in the absence of recombination and assuming that multiple mutations play a minor role, the haplotypic diversity of the sample (defined as the number of haplotypes [34]) should be equal to the number of these SNVs plus one, but it turns out to be systematically higher. In fact, for the swarms infecting the two individuals culled at 35 dpi, recombination could account for about 31% and 28% of the haplotypic diversity in VP1, respectively. In the animal culled after one year of persistent infection, the haplotypic diversity in VP1 is close to saturation (i.e. almost all sequences are different haplotypes) and 75% of it could be attributed to recombination. This rise in the role of recombination in time is consistent with the observed increase in recombination between swarms and supports the persistent replication of the virus even in the carrier state.

#### Reduction in the fitness of recombinants during co-infections

The dynamics of recombination is different when an animal is co-infected by multiple strains, as in our experiment. As discussed before, combinations of alleles belonging to the same strain often have higher fitness even within host, since these combinations have already co-evolved through a range of selective pressures for infectivity and stability and are already adapted. This is a case of positive epistasis between these variants.

Recombination causes the disruption of these beneficial coevolved genetic interactions [35,36]. Hence, the within-host selective pressures due to epistatic interactions tend to reduce the number of recombinants and therefore the effective rate of recombination [27]. This effect is visible even in our data. From Figure 5, the number of recombinants with two VP1 blocks of different origin is suppressed by an order of magnitude with respect to the naive expectation based on the inferred rates. From a back-of-the-envelope computation, the suppression factor can be estimated as *e*^−*s*·*t*^ where *s* is the fitness disadvantage of recombinants within the host and *t* is the number of viral generations since the beginning of the infection. Since a replication cycle of FMDV is completed in about 4 hours [37], we estimate that intra-host epistatic interactions in VP1 are quite strong, with selection coefficients up to *s* ≈ 0.1 per generation.

Given the further selective constraints on infectivity and transmissibility, recombination between different strains could easily generate recombinants with suboptimal combinations of variants not only for within-host growth, but for between-host transmission as well; that would then significantly reduce the probability of observing these recombinants in other hosts. This effect increases with the amount and strength of epistatic interactions, but it occurs even for weak epistasis among many variants [30].

#### The interplay of recombination and epistasis results in the exchange of short sequences

At high recombination rates, there would be a number of recombinants that are almost identical to one of the original strains but for short sequence stretches coming from the other strain. The role of recombination is to mediate these exchanges of short fragments that tend to have a limited impact on fitness and could even form new beneficial allelic combinations. Sequences derived from such exchanges may be transmitted and infect other animals, hence playing a role in the generation of capsid genetic diversity at epidemiological scales. This phenomenon is illustrated in Fig 7B. These exchanges of short recombinant fragments would be almost undetectable from phylogenetic analyses since they would likely be attributed to convergent point mutations, but would actually be originating from co-infection and recombination.

Some indirect evidence for this phenomenon can be found in the samples obtained via tonsil swabs and probangs from four buffaloes involved in the experiment, all of them infected with the same inoculum. These samples show almost no internal variability, while at the consensus level, their sequences correspond to recombinants of the two initial swarms (Cortey et al, companion paper). In these samples, we actually observe such exchanges of short fragments, most of them being about a few tens of bases long. In fact, these recombinant sequences show a clear asymmetry in the amount of bases derived from each swarm: the average contribution of the major swarm of the inoculum is less than 20% and it is scattered in small fragments with a median estimated length of 27 bases, much smaller than the median length of ~ 130 expected for randomly located recombination breakpoints (*p* < 10^−8^ by Mann-Whitney U-test). Each of these fragments contains on average 1-2 swarm-specific variants.

This recombination-mediated exchange of short fragments is a potentially relevant mechanism for genetic exchange between capsid sequences of different FMDV strains. Its evolutionary role could mimic what occurs in non-structural proteins, where the exchange of large fragments and the “mosaic” structure of the genome play an important role in long-term viral evolution by spreading genetic variability across different serotypes.

#### Mismatch between intra- and inter-host recombination rates

Within-host recombination between different FMDV strains during co-infections sometimes results in sequences of high fitness containing large recombinant fragments, which can be transmitted and are able infect other animals, hence playing a role in the long-term evolution of the virus.

These recombination events can also be inferred at phylogenetic scales, i.e. from FMDV sequences collected from different animals and locations. In fact, in the presence of recombination different regions of the genome might have different genealogical trees. If the recombinant fragments are large enough, this signal can be detected in phylogenetic analyses. Such analyses were performed in the past to infer FMDV recombination breakpoints from phylogenetic incompatibilities [18,19].

There is a clear mismatch between the within-host recombination rates observed in our experiment and the much lower recombination rates inferred from phylogenetic analyses. As an example, the number of recombination events in the capsid region inferred in [18] for the whole FMDV phylogeny is similar to the number of intra-host events that we observe after one year of persistent infection in a single individual! In addition, while our findings imply that structural proteins recombine less than non-structural ones located in flanking regions in the genome, this difference in recombination rates appears to be much less extreme than the one observed on a phylogenetic scale. Genome-wide differences in the patterns of within-host versus phylogenetic recombination have been studied in a companion paper [38], yielding qualitatively similar results.

However, there is another key difference between this study and previous studies. Previous phylogenetic investigations focused on recombination between highly divergent sequences: in [19] only inter-serotype recombination events were considered, while in [18], parent sequences belonged to different serotypes in about 70% of the detected events. In contrast, the two main swarms studied here are very similar and belong to the same topotype. This suggests that divergence-dependent effects that suppress recombination on broad scales could be responsible for the mismatch.

Artefacts like biased detection in phylogenetic analyses could partially explain this result. Inference of recombination by phylogenetic methods depends on the resolution of the tree and the similarity of recombining sequences. Inferring recombination between similar sequences is very difficult, since the trees generated by these events are very similar to each other [39] and there are not enough mutations to resolve the recombining branches. In particular, recombination between very close sequences in the phylogenetic tree is hardly detectable. This affects structural and non-structural proteins in a similar way, since it depends only on the local molecular clock. Hence, this cannot be the only reason for the mismatch.

#### Cross-immunity, population structure and epistasis suppress recombination

There are several other genetic and epidemiological factors that can suppress FMDV recombination in endemic and epidemic contexts. Some of these factors could have a stronger impact on structural proteins than on non-structural ones. The mismatch between intra-host and phylogenetic recombination rates offers new opportunities to study these factors.

One of these factors is cross-immunity. The effective rate of recombination is proportional to the rate of co-infections, since co-infection of the same animal/cell by two different strains is a necessary condition for recombination to occur via template switching [20]. The probability of co-infections depends on the ability of the second strain to escape the immune response induced by the first strain, i.e. on the cross-immunity between strains. Cross-immunity depends mostly on capsid proteins [40] (since they are exposed to the immune system) and it tends to decrease with increasing divergence between the capsid sequences of the two strains, hence suppressing recombination between closely related strains only.

Another related factor is the co-circulation of lineages. Geographical separation of lineages could reduce the probability of co-infection and recombination, since the spatial co-existence of different FMDV lineages in the same area is a prerequisite for non-trivial recombination [20]. However, spatial patterns of FMDV are complex and depend on the endemic/epidemic system considered, so the importance and the actual role of this effect is difficult to estimate.

Finally, selection for infectivity and transmissibility would enhance the role of epistatic genetic interactions between variants of well-adapted strains. Such interactions have been recently detected in phylogenetic studies of influenza A [41,42]. While epistatic interactions in infectivity and between-host transmissibility could be present among all FMDV protein-coding regions, it is reasonable to expect stronger selection in the capsid, due to the amount of structural interactions between capsid proteins and the interplay between opposite selective pressures from stability and from the immune system of the host.

The pattern of suppression of recombination due to these constraints is illustrated in Fig 7C using a simple model of cross-immunity and epistatic incompatibilities. For most reasonable values of cross-immunity and epistatic parameters, the model predicts a strong suppression of the recombination rate in structural proteins. The suppression can be of several orders of magnitude, which would explain the near absence of recombination events in capsid on a phylogenetic scale.

#### Recombination, epistasis and speciation in FMDV

Interestingly enough, it was suggested in [41] that the suppression of recombination due to epistatic interactions could act as a mechanism for viral speciation, playing a similar role as hybrid incompatibilities in the classical Dobzhansky-Muller model of speciation [43, 44]. Sympatric speciation is known to occur in viruses [45, 46] and could be caused precisely by the dependence of epistasis and suppression on the amount of divergence between sequences. In the context of FMDV and other picornaviruses, speciation appears to be inhibited by capsid-swapping [18] and frequent recombination events in the genomic regions coding for non-structural proteins. However, epistatic interactions could lead to a genetic separation between incompatible capsid sequences, being therefore the causal factor in the emergence of different serotypes.

## CONCLUSIONS

In this paper we present the first inference of within-host recombination rates for structural proteins of FMDV. This study is possible thanks to a co-infection of two SAT1 viruses, creating two co-occurrent swarms with a small sequence divergence of about 3% inside the buffalo hosts. The recombination rates during the acute and persistent phases of the infection are about *r*_acute_ ~ 0.2/site/y and *r*_persistent_ ~ 0.005/site/y. This shows that intra-host recombination rates are high and even comparable to the mutation rate. It also suggests that the virus keeps replicating in tonsils and other tissues during the carrier phase, although at a rate 40 times slower than in the acute phase of the infection. We provide high-resolution maps of recombination at the scale of the capsid-coding and flanking regions, showing that recombination is a pervasive phenomenon in the FMDV genome. We also discover a modular structure in the VP1-coding region, formed by two strongly linked genomic blocks. Linkage within and between blocks is maintained by widespread epistatic interactions between beneficial combinations of co-evolved variants as well. These selective pressures act both at the protein and the RNA level. The strength of these epistatic selection coefficients is of order 10^−2^.

Our results suggest that recombination and epistasis play an important and unappreciated role in the evolution of FMDV. Within-host recombination is likely to give a strong contribution to the creation of intra-host diversity in FMDV swarms/quasi-species, reducing the mutational load. During co-infections, it could also transfer genetic diversity from one strain to the other via recombination-mediated exchange of short RNA fragments, hence contributing to the between-host evolution of FMDV sequences and the genetic diversity of the virus at broader scales. However, pervasive epistatic interactions between co-evolved variants would prevent the spread of viruses with recombinant capsid sequences. These interactions might be the key factor for “speciation” of FMDV serotypes at the capsid level.

## Materials and methods

### Sequencing

Details on the experimental setup can be found in [2]. The samples included micro-dissected tissues (pharyngeal tonsil, palatine tonsils and dorsal soft palate) from three buffaloes obtained at 35 or 400 days post-infection with FMDV, which were sequenced by Sanger technology. RNA was extracted from LMD material using RNeasy Micro kit (Qiagen), followed by cDNA synthesis using TaqMan RT reagents (Agilent) and random hexamers, then the VP1 region of SAT1 was amplified using Platinum Taq Hi-Fidelity (Invitrogen) and the following primer pair: 5’-AGTGCTGGACCCGACTTCGA-3’ and 5’-TGTAGCGATCCTTGCCACCGT-3’ and the VP1 fragment was cloned into a TOPO^®^TA vector (Life Technologies). After colony picking and plasmid purification, the fragments were Sanger sequenced using BigDye terminator v3.1 (Applied Biosystems) and M13 primers. More details on these sequences will be provided in a separate publication (Cortey et al, companion paper).

The inoculum used for the experiment, as well as six further samples obtained from tonsil swabs and probangs of four animals culled between 200 and 400 days post inoculation, were sequenced at high throughput. RNA was extracted using RNeasy mini Kit (Qiagen), followed by cDNA synthesis using SuperScript™ III First-Strand Synthesis System (Life Technologies), amplification of the capsid region using Platinum Taq Hi-Fidelity (Invitrogen) and the primer pair 1A1F/2B2R (sequences available on request). Libraries were constructed using Nextera XT DNA Sample Preparation Kit (Illumina) and deep sequenced on a MiSeq system using 300 cycle version 2 reagent cartridges (Illumina) to produce paired end reads of approximately 150 bp each.

A reference sequence for the inoculum was assembled using a sensitive in-house pipeline (Ribeca et al, in preparation) based on SPAdes [48] and additional bespoke software. Reads were aligned to this sequence using the GEM mapper [49] version 3. For the inoculum, reads were mapped to the assembly with a mean read depth of about 30000. Multiple alignment of the assembled sequence of the inoculum with the sequences obtained by Sanger technology was performed by Clustal Omega [50]. All genome positions given in the text are relative to the sequence of SAT1/RV/11/37, which is the prototype of SAT1 viruses. The sequenced region comprises the 3’ end of Lpro, the capsid-coding region as well as 2A, 2B and the 5’ end of 2C, and it aligns to the genomic region SAT1:1562-4579.

### Multi-swarm structure

SNV variants in the inoculum were called by a in-house pipeline using an approximation of the Bayesian calling algorithm in Snape-pooled [51] suitable for high coverage. We considered biallelic variants only and selected the SNVs with *p*-value < 0.05. The sequence of SAT1/KNP/196/91 was used to infer the ancestral allele for each SNV. The derived frequency distribution in the inoculum is clearly bimodal with a gap between 0. 15 and 0.20 (Figure 8A). That makes it easy to separate all SNVs in two classes: common nucleotide variants (0.20 < *f* < 0.55) and the low-frequency variants (*f* < 0.15). The first class contains the variants that differentiate between the swarms, while the second is the internal variability of the two swarms.

**FIG 8.**
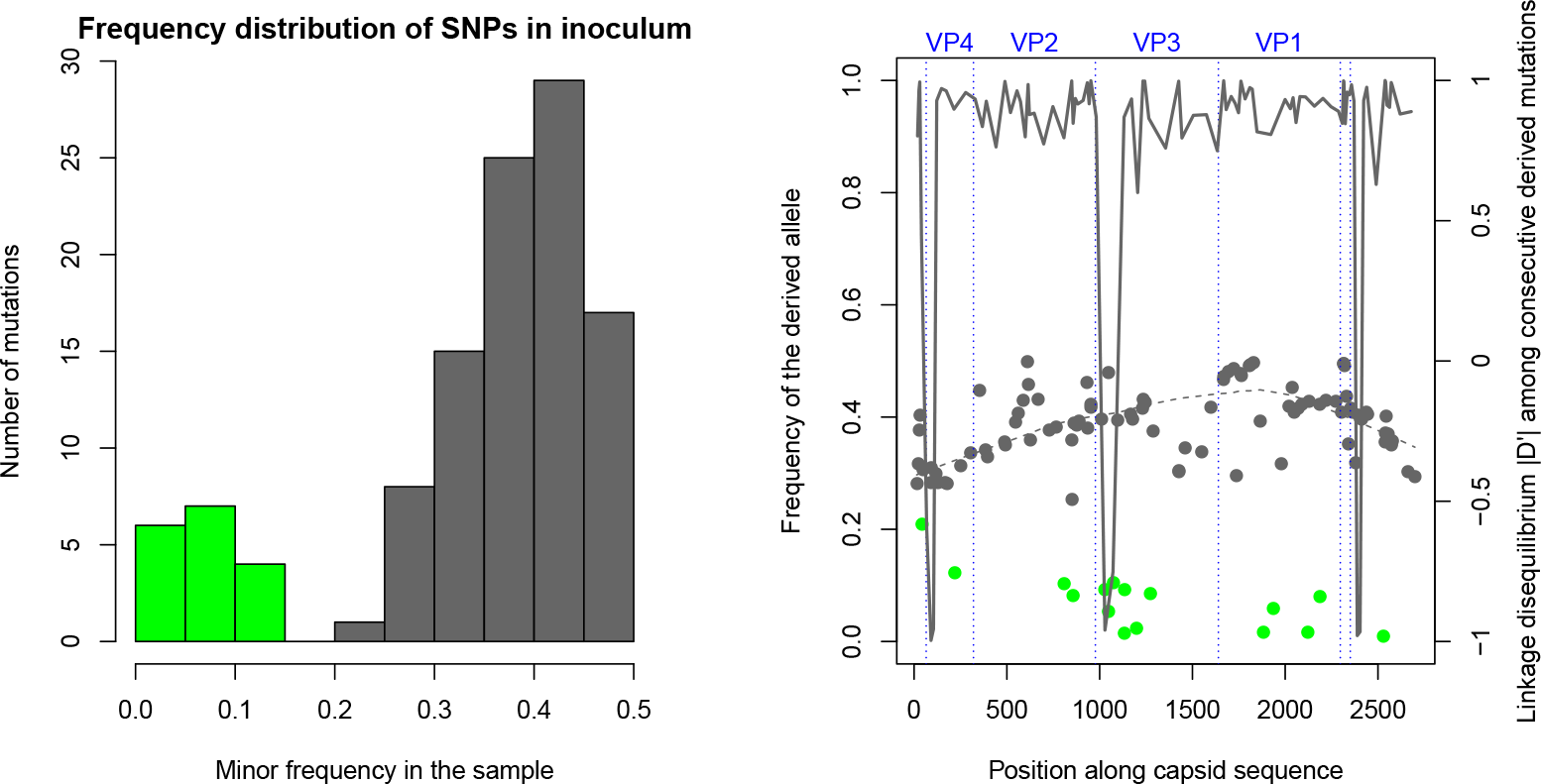
A: Distribution of minor SNV frequencies in the reads from the inoculum. The distribution is clearly bimodal, separating inter-swarm SNVs with minor frequency > 0.2 (grey) from intra-swarm ones (green). B: Location and frequency of the derived allele of SNVs, both inter-swarm (grey points) and intra-swarm (green points). The dashed line is the LOWESS of inter-swarm SNVs. The continuous line shows the linkage disequilibrium *D*′ between pairs of consecutive derived variants.

We estimated the linkage disequilibrium (LD) by the normalised measure *D*′ among all pairs of common variants covered by at least 10^4^ reads. This measure is defined as *D*′ = *D*/*D*_max_ if *D* > 0 and *D*′ = *D*/|*D*_min_| if *D* < 0, where *D* is the classical linkage disequilibrium *D* = *f* (*A*_1_*A*_2_) − *f*(*A*_1_)*f*(*A*_2_) for two SNVs with ancestral alleles *A*_1_ and *A*_2_, while *D*_max_ and *D*_min_ are its maximum and minimum possible value given the frequencies of the variants at the two SNVs [26].

The local haplotype structure of the multi-swarm, with two haplotypes containing ancestral and derived SNV alleles, is clearly illustrated by the concentration of allele frequencies around a value of 0.4 (Figure 8B) and by the high LD between consecutive common variants. In fact, almost all values of |*D*′| are between 0.75 and 1 (Figure 8B). Note that a few mutations have *D*′ ≈ −1, suggesting an erroneous inference of their ancestral state.

We also extracted all nucleotide variants among Sanger sequences and filtered out unreliable SNVs by fitting an empirical model of sequencing errors to our data (see Supplementary Information). After filtering, the VP1-coding sequences from micro-dissections of buffalo tissues show a similar pattern of genetic diversity and LD, already apparent in Figures 1 and 5, although with a larger fraction of intra-swarm variants.

### Evidence of within-host recombination

The main evidence of recombination in the capsid region comes from the LD data from the inoculum in Figure S2. There are clearly many pairs of mutations with low linkage disequilibrium (− 1 ≪ *D*′ ≪ 1), which is a characteristic signature of recombination. Low values of LD can be due to recombination or other spurious factors: (i) sequencing errors; (ii) chimeric reads or similar artefacts of preparation and sequencing protocols; (iii) multiple mutations/backmutations in mutation hotspots. However, none of these factors except recombination can explain the patterns in our data:

- Sequencing errors cannot explain the fact that most putative recombinants contain precisely the same alleles as the two major quasi-species, unless the error rates at each SNV location are extremely skewed towards the same pair of alleles already present. Such an extreme bias seems highly unlikely. Even assuming a moderate bias in error rates, if LD would be caused by sequencing errors, they would be expected to contribute a number of other variants with frequencies similar to the minimum frequency among the four possible pairs of alleles. Instead, Figure 9A shows that the actual contribution of sequencing errors is negligible compared to the predicted one needed to explain our data. A similar argument suggests that mutation hotspots represent an unlikely explanation for the data unless all mutations in these hotspots show a very strong bias towards the two alleles observed at these sites (i.e. most mutations at both sides are back-and-forth mutations between the two alleles).
- Mutation hotspots and sequencing errors cannot explain a pattern of decay of LD with distance along the genome, since they would occur at each site independently, no matter the location. But a clear decay of LD with distance is precisely what is observed in the data (see Figure 9B for the inoculum, and Figure 5 for sequences from buffalo tissues), ruling out these explanations. More quantitatively, recombination implies that the mean LD between positions *i* and *j* satisfies 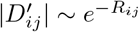 where 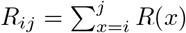 in terms of the cumulative recombination rate per base *R*(*x*). This implies the approximate prediction 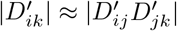 for *i* < *j* < *k*. The results and the predictions for next-to-nearest and next-next-to-nearest SNVs are shown in Figures 9C and 9D respectively. As for the decay of LD, these patterns cannot be replicated by sequencing errors or by back-and-forth mutations.
- The decay of LD is found both in Sanger- and high-throughput-sequenced samples. It is extremely unlikely that these different protocols would generate chimeric reads with similar profiles. Furthermore, chimeric reads and sequencing errors could not cause the decrease in LD (i.e. the increase in cumulative recombination) over time in Figure 2, since they do not depend on time points but on protocols only.
- Finally, for the buffalo culled at 400 dpi, we are able to compare samples from micro-dissections and from a tonsil swab. Deep sequencing of the tonsil swab shows little internal variability. At the consensus level, the sequence of the swab is a complex recombinant of the two initial swarms, with the sequence of 1A-1B (VP4-2) mostly derived from the major swarm in the inoculum and 1C-1D (VP3-1) from the minor one. This consensus-level evidence rules out chimeric sequences or sequencing errors. Tonsil swabs and probangs from other animals also present similar features, although with a reduced contribution of the major swarm of the inoculum.

**Fig 9.**
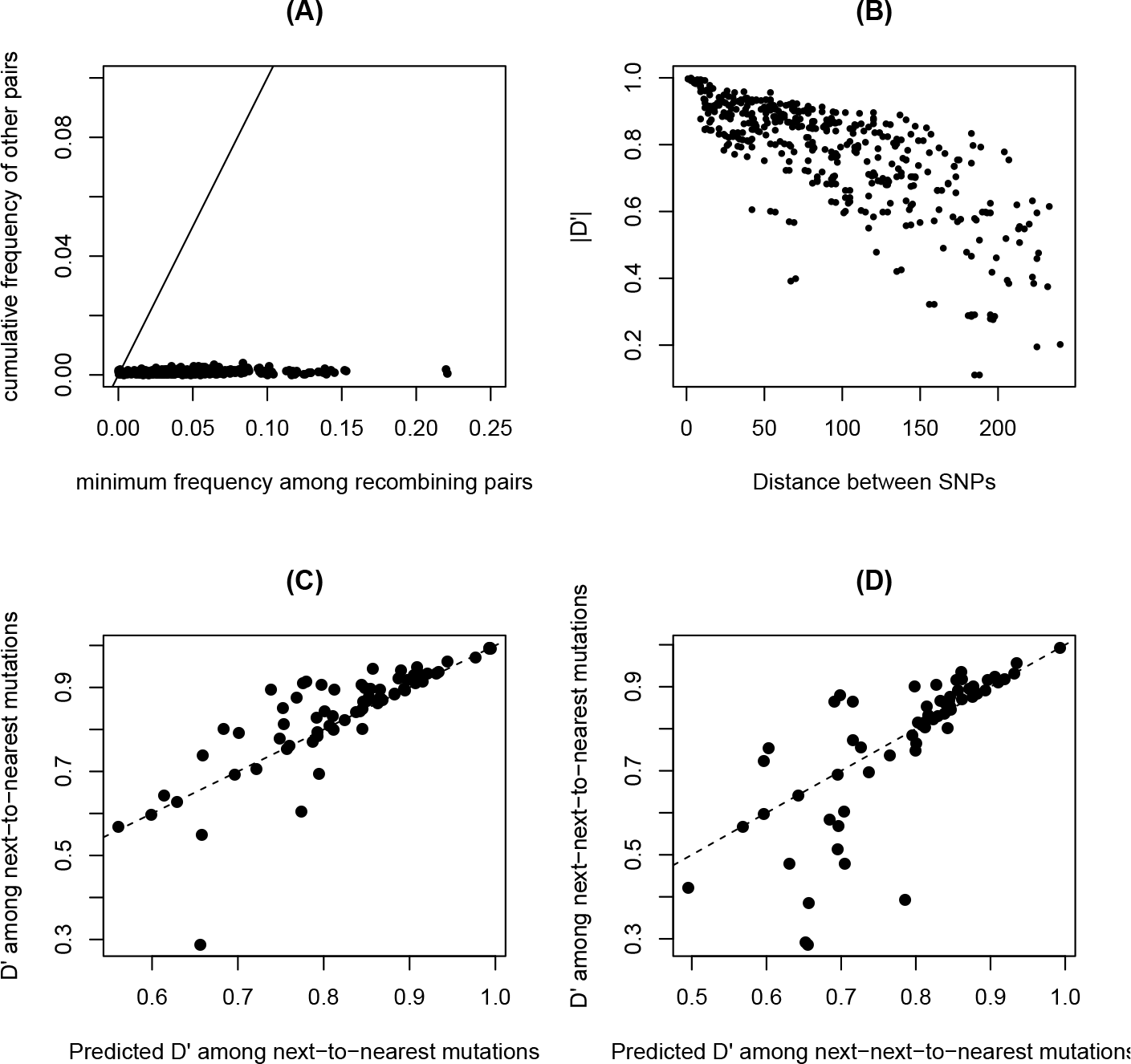
A: Frequency of the least frequent pair of recombinant alleles *f*_*r*,min_ versus the cumulative frequency *f*_*o*_ of all other pairs. For example, if the two main alleles are (C,T) in the first site and (A,G) in the second, then the recombinants are CA, TG, CG, TA; if *f*_CA_ = 0.5, *f*_TG_ = 0.2, *f*_CG_ = 0.1, *f*_TA_ = 0.15, *f*_CC_ = 0.03, *f*_TC_ = 0.02, then *f*_*r*,min_ = 0.1 and *f*_*o*_ = 0.05. Data points are shown for all pairs of polymorphic alleles of frequency > 0.25 covered by at least 10^4^ reads. The black line corresponds to the case of equal frequency. B: Decay of LD measure |*D*′| with distance between SNVs. Data points are shown for all pairs of polymorphic alleles of frequency > 0.25 covered by at least 10^4^ reads. C: *D*′ among next-to-nearest SNVs versus the predicted value from 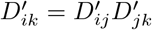. The dashed line corresponds to equality between predicted and estimated value. D: Same but for next-next-to-nearest SNVs.

### Linkage disequilibrium and recombination rates

The high levels of positive LD between all the SNVs of the swarms supports a recent origin for the mixture of swarms. Based on this, we can infer the cumulative rate of recombination since the origin of the swarms using the classical equations for the decay of LD with time: 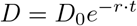, where *r* is the recombination rate per generation and *t* is the time in number of viral generations [26]. The cumulative recombination rate *R* = *r* · *t* for the genomic region between two variants can then be inferred as

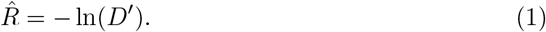

We apply two different statistical approaches to infer the recombination rate for each variant-free interval. The first (“local” approach) is based on the above estimate (1) for consecutive variants only. The second (“global” approach) is the weighted least squares [52] estimate *R*^wls^ from all variants, defined by the equations:

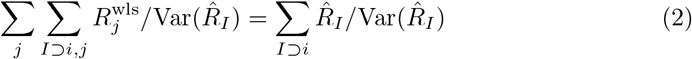

where *i*, *j* denote intervals between consecutive variants and *I* intervals between any pair of variants. We use the approximate form for the variance: 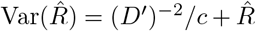, where *c* is the number of reads covering both variants in the pair; the first term comes from the delta method applied to the variance of binomial sampling of *c* sequences (assuming low recombination and similar frequencies for all SNVs), the second from the Poisson noise of the random recombination events. To get comparable results between Sanger and short-reads data, only intervals of length less than 200 bases are used for the “global’ estimate for analyses involving both approaches.

Data from Sanger sequencing of viruses from micro-dissections reveals only weak differentiation between tissues from the same animal. The average estimate of 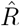 across tissues and the joint estimate from all tissues differ by less than 10%, hence we neglect differences across tissues and compute *D*′ from the pooled set of all sequences from a given animal.

### Epistasis

The cumulative recombination rate between the *i*th and *j*th variant is inferred according to equation (1). The local prediction for the rate is the sum of all rates for the sequence between the *i*th and *j*th variant 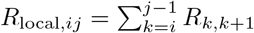. We also implement a non-local prediction defined as 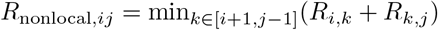. This prediction takes into account the suppression of recombination caused by the the most linked variant in-between, hence *R*/*R*_nonlocal_ provides a better estimate of the epistatic suppression due to the pairwise interaction between the *i*th and *j*th variant alone.

Epistatic selection coefficients *s* are inferred from the solution of

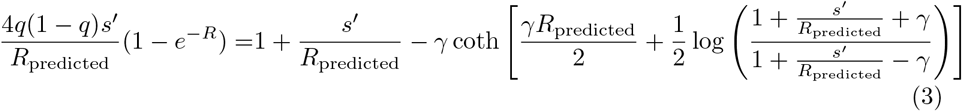

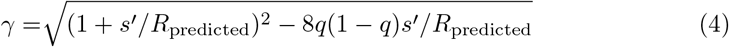

where *s’* = *s* · *t* and *t* is the time in generation since the beginning of the experiment, *R* is the estimate 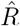 of the cumulative recombination rate between the mutations and *R*_predicted_ is either the corresponding *R*_local_ or *R*_nonlocal_ (see Supplementary Information for details).

## Supporting information

## Acknowledgments

We thank colleagues in the WRLFMD (especially Nick Knowles and Antonello Di Nardo) for useful discussions. The Pirbright Institute receives grant-aided support from the Biotechnology and Biological Sciences Research Council of the United Kingdom (projects BB/E/I/00007035, BB/E/I/00007036 and BBS/E/I/00007039 and grant BB/L011085/1 as part of the joint USDA-NSF-NIH-BBSRC Ecology and Evolution of Infectious Diseases program).

## Supporting information

**S1 File. Supplementary Methods and Figures**. Supplementary Information containing further details on data analysis.

## References

1. Alexandersen S, Zhang Z, Donaldson A, Garland A. The pathogenesis and diagnosis of foot-and-mouth disease. Journal of comparative pathology. 2003;129(1):1–36.

2. Maree F, de Klerk-Lorist LM, Gubbins S, Zhang F, Seago J, Pérez-Martín E, et al. Differential persistence of foot-and-mouth disease virus in African buffalo is related to virus virulence. Journal of virology. 2016;90(10):5132–5140.

3. Mason PW, Grubman MJ, Baxt B. Molecular basis of pathogenesis of FMDV. Virus research. 2003;91(1):9–32.

4. Belsham GJ, Charleston B, Jackson T, Paton DJ. Foot-and-Mouth Disease. eLS. 2009;.

5. Domingo E, Holland J. RNA virus mutations and fitness for survival. Annual Reviews in Microbiology. 1997;51(1):151–178.

6. Knowles N, Samuel A. Molecular epidemiology of foot-and-mouth disease virus. Virus research. 2003;91(1):65–80.

7. Paton DJ, Sumption KJ, Charleston B. Options for control of foot-and-mouth disease: knowledge, capability and policy. Philosophical Transactions of the Royal Society B: Biological Sciences. 2009;364(1530):2657–2667.

8. Gebauer F, De La Torre J, Gomes I, Mateu M, Barahona H, Tiraboschi B, et al. Rapid selection of genetic and antigenic variants of foot-and-mouth disease virus during persistence in cattle. Journal of virology. 1988;62(6):2041–2049.

9. Lauring AS, Andino R. Quasispecies theory and the behavior of RNA viruses. PLoS pathogens. 2010;6(7):e1001005.

10. Domingo E, Sheldon J, Perales C. Viral quasispecies evolution. Microbiology and Molecular Biology Reviews. 2012;76(2):159–216.

11. King AM. Genetic recombination in positive strand RNA viruses. In: RNA Genetics, Volume II, Retroviruses, viroids, and RNA recombination. CRC Press Albany, NY; 1988. p. 149–165.

12. Carrillo C, Tulman E, Delhon G, Lu Z, Carreno A, Vagnozzi A, et al. Comparative genomics of foot-and-mouth disease virus. Journal of virology. 2005;79(10):6487–6504.

13. Lewis-Rogers N, McClellan DA, Crandall KA. The evolution of foot-and-mouth disease virus: impacts of recombination and selection. Infection, Genetics and Evolution. 2008;8(6):786–798.

14. McCahon D, Slade W, Priston R, Lake J. An extended genetic recombination map for foot-and-mouth disease virus. Journal of General Virology. 1977;35(3):555–565.

15. King A, Slade W, Newman J, McCahon D. Temperature-sensitive mutants of foot-and-mouth disease virus with altered structural polypeptides. II. Comparison of recombination and biochemical maps. Journal of virology. 1980;34(1):67–72.

16. Tosh C, Hemadri D, Sanyal A. Evidence of recombination in the capsid-coding region of type A foot-and-mouth disease virus. Journal of general virology. 2002;83(10):2455–2460.

17. Tosh C, Sanyal A, Hemadri D. Genetic and antigenic analysis of a recombinant foot-and-mouth disease virus. Current Science. 2002; p. 1016–1019.

18. Heath L, Van Der Walt E, Varsani A, Martin DP. Recombination patterns in aphthoviruses mirror those found in other picornaviruses. Journal of Virology. 2006;80(23):11827–11832.

19. Jackson A, O’neill H, Maree F, Blignaut B, Carrillo C, Rodriguez L, et al. Mosaic structure of foot-and-mouth disease virus genomes. Journal of General Virology. 2007;88(2):487–492.

20. Worobey M, Holmes EC. Evolutionary aspects of recombination in RNA viruses. Journal of General Virology. 1999;80(10):2535–2543.

21. Shriner D, Rodrigo AG, Nickle DC, Mullins JI. Pervasive genomic recombination of HIV-1 in vivo. Genetics. 2004;167(4):1573–1583.

22. Froissart R, Roze D, Uzest M, Galibert L, Blanc S, Michalakis Y. Recombination every day: abundant recombination in a virus during a single multi-cellular host infection. PLoS biology. 2005;3(3):e89.

23. Neher RA, Leitner T. Recombination rate and selection strength in HIV intra-patient evolution. PLoS computational biology. 2010;6(1):e1000660.

24. Batorsky R, Kearney MF, Palmer SE, Maldarelli F, Rouzine IM, Coffin JM. Estimate of effective recombination rate and average selection coefficient for HIV in chronic infection. Proceedings of the National Academy of Sciences. 2011;108(14):5661–5666.

25. Paton DJ, Gubbins S, King DP. Understanding the transmission of foot-and-mouth disease virus at different scales. Current opinion in virology. 2018;28:85–91.

26. Hartl DL, Clark AG, Clark AG. Principles of population genetics. vol. 116. Sinauer associates Sunderland; 1997.

27. Franklin I, Lewontin R. Is the gene the unit of selection? Genetics. 1970;65(4):707–734.

28. Cottam EM, Haydon DT, Paton DJ, Gloster J, Wilesmith JW, Ferris NP, et al. Molecular epidemiology of the foot-and-mouth disease virus outbreak in the United Kingdom in 2001. Journal of Virology. 2006;80(22):11274–11282.

29. Wright CF, Knowles NJ, Di Nardo A, Paton DJ, Haydon DT, King DP. Reconstructing the origin and transmission dynamics of the 1967-68 foot-and-mouth disease epidemic in the United Kingdom. Infection, Genetics and Evolution. 2013;20:230–238.

30. Neher RA, Shraiman BI. Competition between recombination and epistasis can cause a transition from allele to genotype selection. Proceedings of the National Academy of Sciences. 2009;106(16):6866–6871.

31. Xiao Y, Dolan PT, Goldstein EF, Li M, Farkov M, Brodsky L, et al. Poliovirus intrahost evolution is required to overcome tissue-specific innate immune responses. Nature communications. 2017;8(1):375.

32. Charpentier C, Nora T, Tenaillon O, Clavel F, Hance AJ. Extensive recombination among human immunodeficiency virus type 1 quasispecies makes an important contribution to viral diversity in individual patients. Journal of virology. 2006;80(5):2472–2482.

33. Simon-Loriere E, Holmes EC. Why do RNA viruses recombine? Nature Reviews Microbiology. 2011;9(8):617.

34. Gregori J, Perales C, Rodriguez-Frias F, Esteban JI, Quer J, Domingo E. Viral quasispecies complexity measures. Virology. 2016;493:227–237.

35. Martin DP, Van der Walt E, Posada D, Rybicki EP. The evolutionary value of recombination is constrained by genome modularity. PLoS genetics. 2005;1(4):e51.

36. Monjane AL, Martin DP, Lakay F, Muhire BM, Pande D, Varsani A, et al. Extensive recombination-induced disruption of genetic interactions is highly deleterious but can be partially reversed by small numbers of secondary recombination events. Journal of virology. 2014;88(14):7843–7851.

37. Han SC, Guo HC, Sun SQ, Jin Y, Wei YQ, Feng X, et al. Productive entry of foot-and-mouth disease virus via macropinocytosis independent of phosphatidylinositol 3-kinase. Scientific reports. 2016;6:19294.

38. Ferretti L, Di AN, Singer B, Lasecka-Dykes L, Logan G, Wright C, et al. Within-Host Recombination in the Foot-and-Mouth Disease Virus Genome. Viruses. 2018;10(5).

39. Ferretti L, Disanto F, Wiehe T. The effect of single recombination events on coalescent tree height and shape. PloS one. 2013;8(4):e60123.

40. Bøtner A, Kakker NK, Barbezange C, Berryman S, Jackson T, Belsham GJ. Capsid proteins from field strains of foot-and-mouth disease virus confer a pathogenic phenotype in cattle on an attenuated, cell-culture-adapted virus. Journal of General Virology. 2011;92(5):1141–1151.

41. Villa M, Lässig M. Fitness cost of reassortment in human influenza. PLoS pathogens. 2017;13(11):e1006685.

42. White MC, Lowen AC. Implications of segment mismatch for influenza A virus evolution. Journal of General Virology. 2017;.

43. Orr HA. The population genetics of speciation: the evolution of hybrid incompatibilities. Genetics. 1995;139(4):1805–1813.

44. Gavrilets S. Hybrid zones with Dobzhansky-type epistatic selection. Evolution. 1997;51(4):1027–1035.

45. Kitchen A, Shackelton LA, Holmes EC. Family level phylogenies reveal modes of macroevolution in RNA viruses. Proceedings of the National Academy of Sciences. 2011;108(1):238–243.

46. Meyer JR, Dobias DT, Medina SJ, Servilio L, Gupta A, Lenski RE. Ecological speciation of bacteriophage lambda in allopatry and sympatry. Science. 2016; p. aai8446.

47. Luksza M, Lässig M. A predictive fitness model for influenza. Nature. 2014;507(7490):57.

48. Bankevich A, Nurk S, Antipov D, Gurevich AA, Dvorkin M, Kulikov AS, et al. SPAdes: a new genome assembly algorithm and its applications to single-cell sequencing. Journal of computational biology. 2012;19(5):455–477.

49. Marco-Sola S, Sammeth M, Guigó R, Ribeca P. The GEM mapper: fast, accurate and versatile alignment by filtration. Nature methods. 2012;9(12):1185.

50. Sievers F, Wilm A, Dineen D, Gibson TJ, Karplus K, Li W, et al. Fast, scalable generation of high-quality protein multiple sequence alignments using Clustal Omega. Molecular systems biology. 2011;7(1):539.

51. Raineri E, Ferretti L, Esteve-Codina A, Nevado B, Heath S, Pérez-Enciso M. SNP calling by sequencing pooled samples. BMC bioinformatics. 2012;13(1):239.

52. Strutz T. Data fitting and uncertainty: A practical introduction to weighted least squares and beyond. Vieweg and Teubner; 2010.

