## Supplementary material for "Pervasive within-host recombination and epistasis as major determinants of the molecular evolution of the Foot-and-Mouth Disease Virus capsid"

#### S1 Genetic content of the inoculum

After the removal of reads from contaminations, all sequenced FMDV reads belong to the SAT1 serotype. The consensus sequence of VP1 is almost identical to the sequence of SAT1/KNP/196/91, but all sequences in the inoculum present also an insertion of 21 bases with respect to SAT1/KNP/196/91. There are 26 SNPs at frequency $> 1\%$ in VP1, of which 22 have an allele at frequency about 0.45. This corresponds to a strong haplotype structure in the inoculum, with at least two main variants (or better, two swarms around these variants, plus their recombinants) differing by about 3% in their VP1 sequence. The majority of these mutations are at the third codon position (17 out of 26, $p = 4 \times 10^{-3}$ by multinomial test), suggesting purifying selection pressure on the virus.

#### S2 Error model for Sanger sequences

All sequences obtained by Sanger sequencing from two animals culled at 35 dpi and one culled at 400 dpi were pooled together and aligned to the sequence of the inoculum using MAFFT-ginsi [**?**]. From this alignment, the distribution of the counts of all minor alleles in the sample was extracted. Since the distribution of sequencing errors was not known, we fitted an error model to the low-frequency part of the allele distribution, under the conservative assumption that all low-frequency alleles were sequencing errors. Simple models of random errors predict a Poisson distribution of

*

error frequencies (resulting from rare, independent errors in each sequence) and more generally the tail of the error distribution is expected to be exponential. In fact the distribution of alleles present in three to ten sequences was well-fitted by a geometric distribution $P(c) = 186 \cdot 0.69^{(c-1)}$ (Figure S1), which is the discrete version of an exponential distribution. From this geometric fit, the number of expected false SNPs in the sample above a threshold count $\bar{c}$ was estimated as $E[\text{false SNPs}] = 186 \cdot 0.69^{\bar{c}}/(1 - 0.69)$. We fix the threshold by requiring that $\mathrm{E}[\#\ \text{false SNPs}] < 0.5$, that implies $\bar{c} = 20$.

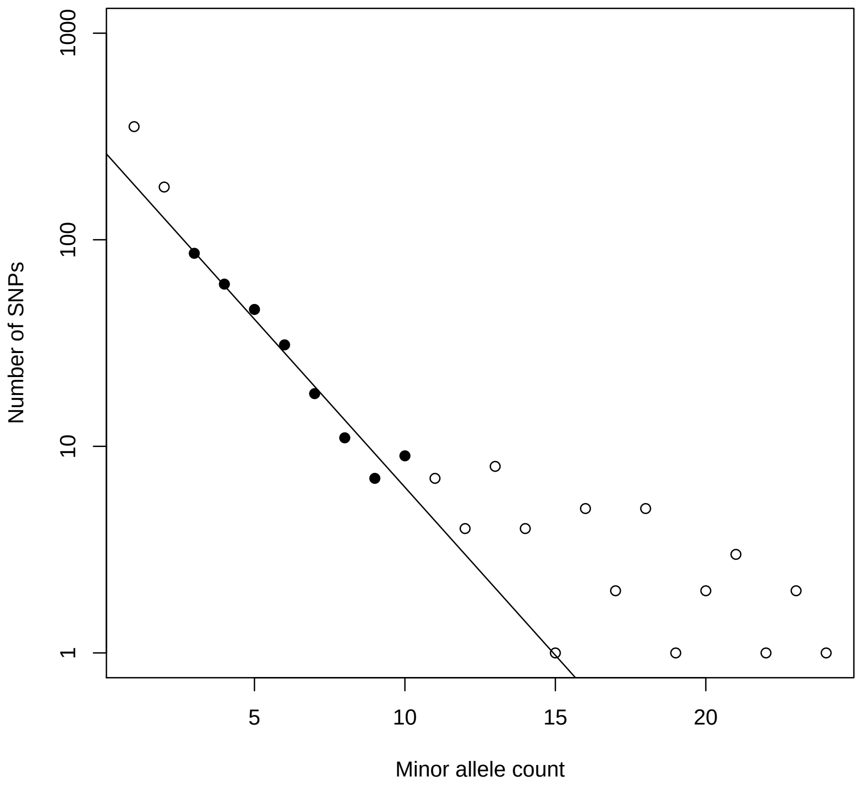


Figure S1: The low-frequency part of the distribution of minor allele counts. The data with counts between 3 and 10 (black dots) were fitted by an exponential model (black line)

### S3 Linkage disequilibrium from short reads

Consider two variables sites in the genome. If there are $c$ reads covering both sites, it is possible to estimate the linkage disequilibrium between their alleles by restricting the analysis on these $c$ reads and computing $D$ and $D'$ from standard expressions for multiple haploid sequences [**?**].

The result of this computation for inter-swarm SNPs in the inoculum is shown in Figure S2.

For pairs of linked loci of similar frequency $q$ and covered by $c$ reads, the sampling error on the estimates of $D'$ [**?**] is about

$$\sigma^2(D') = \frac{(1 - D')(1 + (\frac{1}{q(1-q)} - 3)D' + 3D'^2)}{c} \tag{S1}$$

hence the delta method implies that the variance of the estimates of $R = -\log(D')$ is about

$$\sigma^2(R) \approx \frac{\sigma^2(D')}{D'^2} = \frac{(1-D')(1+(\frac{1}{q(1-q)}-3)D'+3D'^2)}{cD'^2} \tag{S2}$$

For values of $R > 0.05$, the relative error $\sigma(R)/R \lesssim 10\%$ is reasonably small. Even for smaller values of $R > 0.01$, the error $\sigma(R)/R \lesssim 25\%$ is under control.

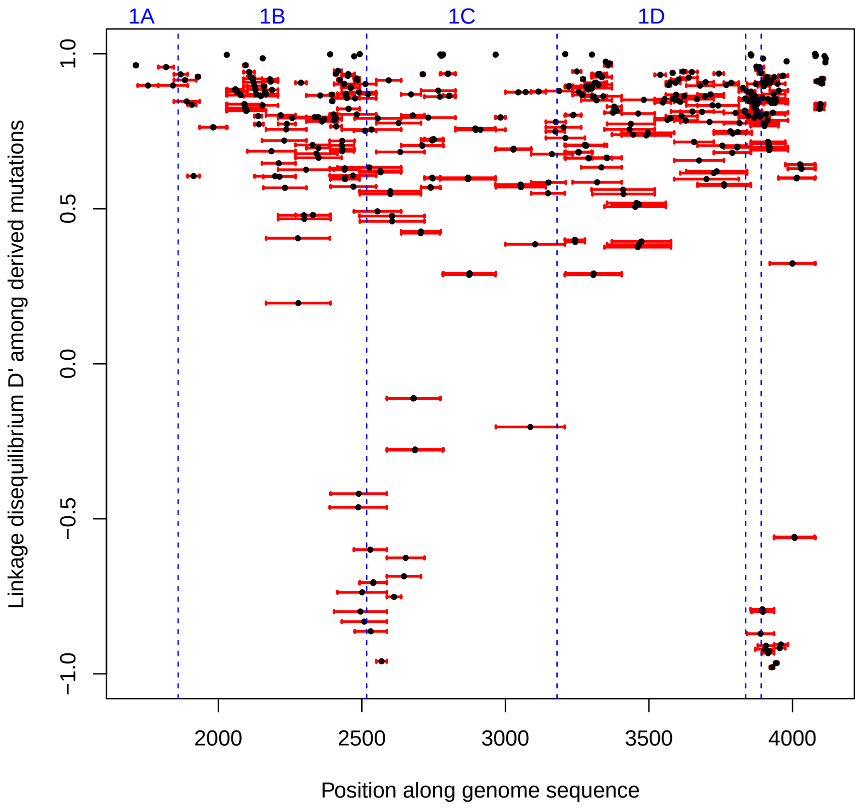


Figure S2: Normalised linkage disequilibrium $D'$ between pairs of derived swarm-specific variants covered by at least $10^4$ reads. Red bars illustrate the interval between variants, with a black dot at the mid-point.

### S4 Multi-swarm structure

The description of the peculiar structure of the quasi-species is based on three pieces of evidence:

- The distribution of minor allele frequencies in the inoculum is strongly bimodal (Figure 8A in the Main Text), with a expected tail of low-frequency alleles disappearing before frequency 0.2, and a large number of intermediate-frequency alleles peaked at 0.4. This already is a strong hint of haplotype structure. The derived alleles show a similar distribution of frequencies and the intermediate-frequency SNPs are distributed uniformly along the sequence (Figure 8B in the Main Text).

- A haplotype structure is defined by a strong linkage disequilibrium (LD) among derived alleles. In the inoculum, LD can be measured only between close SNPs. Figure XXXB in the Main Text shows that the LD between the derived alleles of consecutive SNPs is very strong ($D' \approx 1$), therefore supporting a local haplotype structure. The few mutations with $D' \approx -1$ point to an erroneous inference of their ancestral state.

- The Sanger sequences from microdissections of infected buffalos were sampled after the acute phase of the infection, hence they are affected by selection and recombination in a complex way. However, when looking at the SNPs already present in the inoculum, they still show a bimodal distribution of frequencies (Figure S3) and signatures of a strong haplotype structure (Figure 1 in the Main Text).

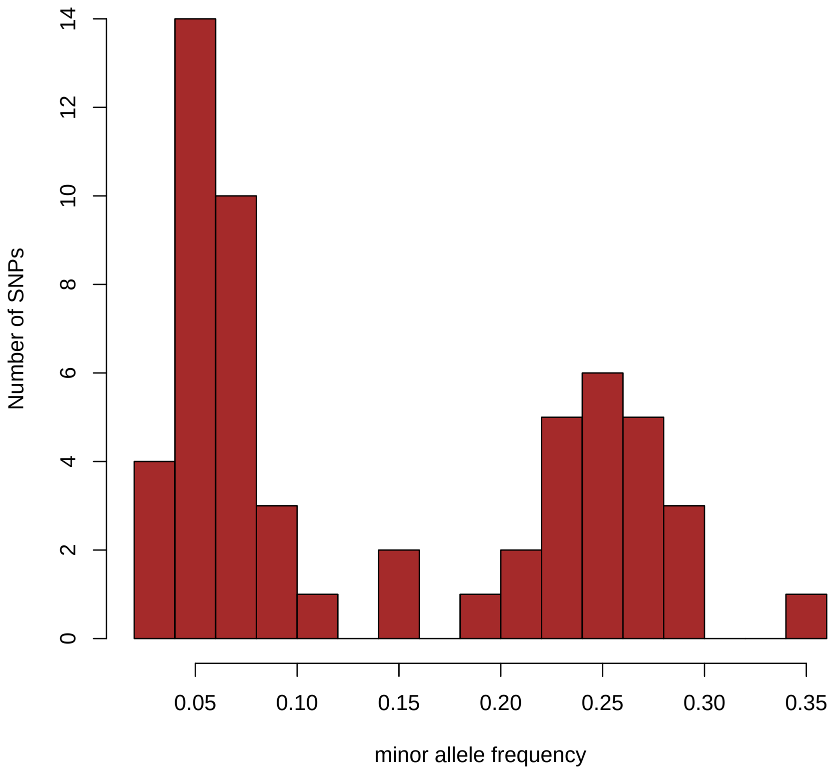


Figure S3: Distribution of SNP frequencies in viral sequences from all micro-dissections of buffalo tissues.

These data support a strong haplotype structure of the quasi-species, with clouds of genotype concentrated around two well-defined haplotypes and their recombinants. The frequency of these haplotypes in VP1 can be estimated by taking the average of the frequencies of the derived and ancestral variants in the SNPs, obtaining 0.44 and 0.56 respectively.

The minor haplotype among the buffalo sequences in Figure 1 does not correspond to any of the two main haplotypes of the quasi-species. It is very similar to the major haplotype of the inoculum, but with two “fixed” variants identical to the minor haplotype. Nevertheless, since it is observed at similar frequencies in the different individuals and tissues, it should have been present in the inoculum: otherwise, if it would have emerged as a random recombinant, its evolutionary outcomes would have been highly stochastic. An estimate of the initial frequency of this haplotype in the inoculum can be obtained by the fraction of reads that (i) cover one of the two sites with “fixed” variants and a neighbouring SNP and (ii) are compatible with this haplotype. The estimated fraction of reads satisfying both conditions (i),(ii) among all reads satisfying condition (i) is about 0.02.

##### S4.1 Viral sequences from micro-dissections

The frequencies of all SNPs among the viral sequences from micro-dissection are shown in Figure S3. Their distribution is again bimodal. New SNPs (corresponding to monomorphic sites in the VP1 sequence of the inoculum) do not show any significant purifying selection ($p > 0.05$ by multinomial test on their codon position).

Two main haplotypes (plus several recombinants) are present among the sequences post-inoculation. The major haplotype is the same as the minor haplotype in the inoculum. Instead, the second main haplotype post-inoculation is not the major haplotype of the inoculum, but it is rather a (possibly recombinant) variant of that haplotype differing only by 0.2% in the VP1 sequence. The VP1 sequences of the minor haplotype in the inoculum and of this recombinant differ in 20 positions out of the 22 SNPs at intermediate frequency in the inoculum, while the remaining two SNPs at intermediate frequency in the inoculum are fixed post-inoculation. Assuming that this variant was already present in the inoculum - which is the most parsimonious hypothesis - it is possible to estimate its initial frequency at about 2% or less in the inoculum, as discussed above. Note that the value of linkage disequilibrium $D$ among the remaining variants is affected by these changes in frequency post inoculation, but its normalized value $D'$ is not.

#### S5 Epistasis

As discussed in the Main text, from the $i$th and $j$th variant ($i < j$) and the inferred recombination rate between them $R_{i,j} = -\log(D'_{i,j})$, we define the predicted recombination rates as

$$R_{local} = \sum_{k=i}^{j-1} R_{k,k+1} \tag{S3}$$

$$R_{nonlocal} = \min_{k \in [i+1, j-1]} (R_{i,k} + R_{k,j}) \tag{S4}$$

The suppression of recombination $R/R_{nonlocal}$ is shown in Figure S4.

The suppression of recombination is related to the selection coefficients, but it is not a direct measure. To infer the actual strength of the epistatic selection coefficients, we need an explicit model relating them to the suppression of recombination.

Consider two sites with variants of fixed frequency $q$, recombining with a recombination rate $r$. Assume also that all recombinants have a fitness disadvantage $-s$. Initially, the two sites are fully linked and $D' = 1$. The expected evolution of recombinants between these sites can be written in terms of a single differential equation for the fraction of recombinants $f_r$:

$$\frac{df_r}{dt} = r(2q(1-q) - f_r) - sf_r(1-f_r) \tag{S5}$$

which can be translated into an equation for the evolution of the linkage disequilibrium $D' = 1 - \frac{f_r}{2q(1-q)}$:

$$\frac{dD'}{dt} = -rD' + s(1-D')[1 - 2q(1-q)(1-D')] \tag{S6}$$

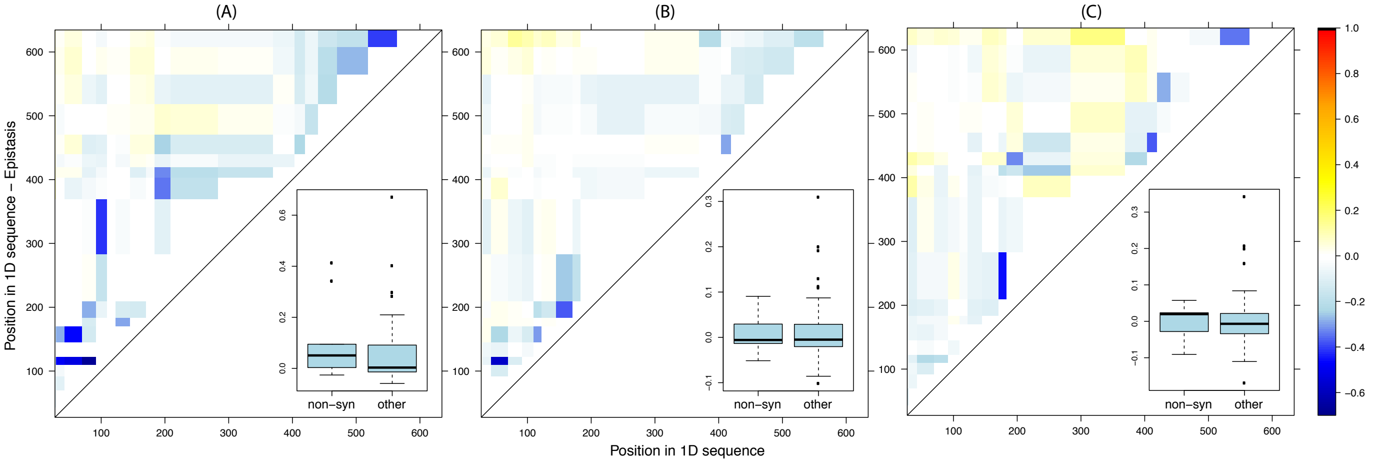


Figure S4: Suppression of recombination $\log_{10}(R/R_{nonlocal})$ corrected to extract only the effect of pairwise interactions. Animals culled at 35 dpi (A,B) and 400 dpi (C). In the insets, boxplots of the strength of epistasis as measured by the inverse suppression $\log_{10}(R_{nonlocal}/R)$ between pairs of non-synonymous mutations and between other pairs.

In turn, this can be solved in terms of the overall effective recombination rate $R = -\log(D')$ as

$$R = -\log\left[1 - \frac{1 + s/r - \gamma\coth\left[\frac{\gamma r t}{2} + \frac{1}{2}\log\left(\frac{1+s/r+\gamma}{1+s/r-\gamma}\right)\right]}{4q(1-q)s/r}\right] \quad \text{(S7)}$$

where $\gamma = \sqrt{(1+s/r)^2 - 8q(1-q)s/r}$.

In this formula, we can replace the recombination rate $r \cdot t \to R_{predicted}$ and the ratio $s/r = \frac{s \cdot t}{r \cdot t} \to s'/R_{predicted}$ and we end up with an implicit equation for the rescaled selection coefficient $s' = s \cdot t$:

$$\frac{4q(1-q)s'}{R_{predicted}}(1 - e^{-R}) = 1 + \frac{s'}{R_{predicted}} - \gamma\coth\left[\frac{\gamma R_{predicted}}{2} + \frac{1}{2}\log\left(\frac{1 + \frac{s'}{R_{predicted}} + \gamma}{1 + \frac{s'}{R_{predicted}} - \gamma}\right)\right] \quad \text{(S8)}$$

$$\gamma = \sqrt{(1 + s'/R_{predicted})^2 - 8q(1-q)s'/R_{predicted}} \quad \text{(S9)}$$

For numerical reasons, we estimate $s' = 0$ if the suppression of recombination is too small, i.e. if $R - R_{predicted} > -0.1$.

The results are shown in Figure S5. The inferred coefficients $s'$ based on $R_{predicted} = R_{local}$ are shown in the top-left triangles of Figure S5(A,B,C,D). Most of the coefficients lie in the range $s' \sim 0 - 5$; if the generation time is about 4 hours, the selective coefficients can be estimated in the range $s \sim 0 - 0.1$. The bottom-right triangles in Figure S5(A,B,C,D) show the inferred coefficients $s'$ corrected to estimate the strength of pairwise interactions, based on $R_{predicted} = R_{nonlocal}$. Most of the corresponding coefficients $s$ lie in the range $s \sim 0 - 0.02$. These coefficients tend to be higher for interactions between non-synonymous mutations, as shown in the boxplots in Figure S5(E,F,G).

To prove the significance of this effect, we compare the median $s'$ from $R_{nonlocal}$ for interactions involving only non-synonymous variants and for interactions involving only synonymous ones via Mann-Whitney U-test. We obtain $p < 0.04$ after combining the $p$-values for each individual by Fisher's method.

The localisation of the five non-synonymous variants in the capsid is illustrated in Figure 6. Only two of the variants are exposed, and only one is in a known epitope. On the other hand, three out of four interacting pairs are localised in close proximity in the FMDV capsid. This suggests that they could be compensatory mutations and that these interactions are probably related to the stability of the VP1 protein and the capsid structure.

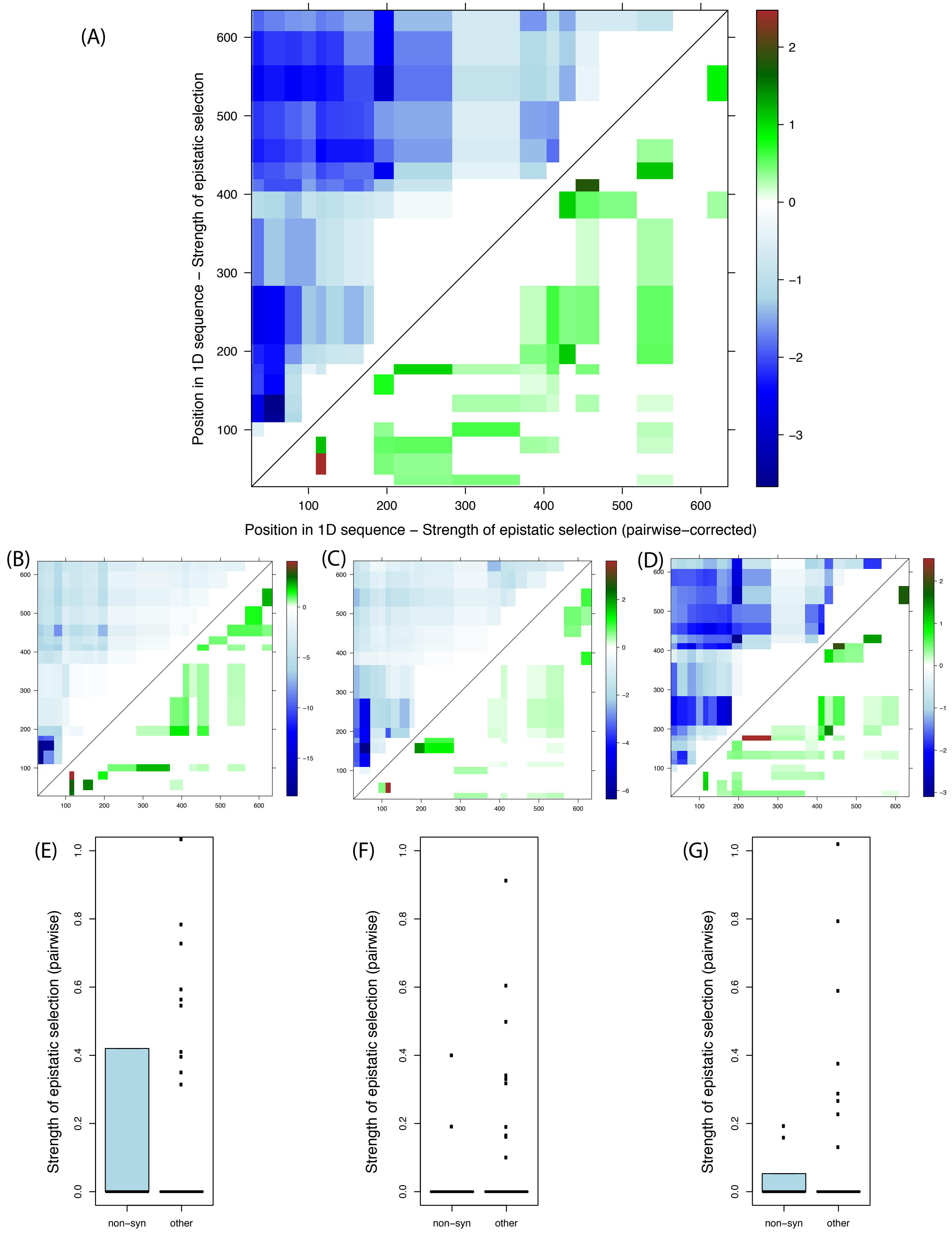


Figure S5: Different measures of the strength of epistatic selection coefficients (top: all, bottom left and centre: buffalo culled at 35 dpi, bottom right: at 400 dpi). See text for explanations.
